# Double Stranded DNA Breaks and Genome Editing Trigger Ribosome Remodeling and Translational Shutdown

**DOI:** 10.1101/486704

**Authors:** Celeste Riepe, Elena Zelin, Stacia K. Wyman, David N. Nguyen, Jin Rui Liang, Phillip A. Frankino, Zuriah A. Meacham, Jonathan T. Vu, Alexander Marson, Nicholas T. Ingolia, Jacob E. Corn

**Affiliations:** Department of Molecular and Cell Biology, University of California, Berkeley, Berkeley, California, 94720, USA; Innovative Genomics Institute, University of California, Berkeley, Berkeley, California, 94704, USA; Department of Microbiology and Immunology, University of California, San Francisco Medical Center, San Francisco, California, 94143, USA; Innovative Genomics Institute, University of California, San Francisco, San Francisco, California, 94143, USA

## Abstract

DNA damage activates a robust transcriptional stress response, but much less is known about how DNA impacts translation. The advent of genome editing via a Cas9-induced DNA double-strand break has intensified interest in understanding cellular responses to DNA damage. Here we find that DNA double-strand breaks (DSBs) induced by Cas9 or other damaging agents lead to a reduction of core ribosomal proteins, RPS27A and RPL40, and that the loss of these proteins is post-transcriptional and p53-independent. DSBs furthermore lead to the shutdown of translation through phosphorylation of eukaryotic initiation factor 2 alpha, and altering these signals affects genome editing outcomes. This DSB translational response is widespread and precedes the transcriptional response. Our results demonstrate that even a single double-strand break can lead to ribosome remodeling and reduced translational output, and suggest caution in interpreting cellular phenotypes measured immediately after genome editing.

## Introduction

Unrepaired DNA damage can lead to lethal mutations and contributes to cancer initiation and progression. Cells have thus evolved a variety of responses to protect their genomes from a myriad of chemical and environmental insults. Double-strand breaks pose a particularly acute danger, as they may cause the wholesale loss of genetic information and require dramatic repair processes. In humans, cells with double-strand breaks arrest until repair is completed and undergo programmed cell death if repair is unsuccessful.

Double-strand breaks provoke a distinctive transcriptional response. Activation of the transcription factor p53 is a hallmark of the DSB response, leading to transcriptional reprogramming, cell cycle arrest, or in cases of severe damage, apoptosis (Joerger and Fersht, 2016). Deficiency in p53 signaling is also pivotal to the progression of many cancers, allowing neoplasms to accumulate DNA damage that results in mutations rapid tumor evolution. In addition to its critical role in maintaining genomic integrity, the cellular response to DSBs is essential to genome editing methods like CRISPR-Cas9. Cas9 editing relies on introducing a targeted double-strand break within a genome, which the cell repairs through error-prone non-homologous end joining (NHEJ) or through templated, homology directed repair (HDR). HDR from even a single Cas9-mediated DSB can induce low levels of p53 signaling, which can have negative consequences for cell fitness and genome editing outcomes (Haapaniemi et al., 2018; Ihry et al., 2018).

Although DSBs are known to initiate transcriptional changes, less is understood about the role of translation in the DNA damage response. A purely transcriptional reaction to a genetic insult leaves a gap in response, potentially exposing a cell to the impact of damaged DNA during a critical time window in which damage had raised an alarm but newly transcribed mRNAs have not accumulated. While transcriptional changes can modulate protein abundance hours or days after a genomic insult, translational control can enact regulatory programs within minutes of an environmental stress (Andreev et al., 2015; Sidrauski et al., 2015).

We thus sought to characterize how cells respond to DNA damage at the translation level, and in particular, how cells respond to a single double-strand break during Cas9-mediated genome editing. We serendipitously found that cells temporarily deplete core ribosomal proteins, RPS27A and RPL40, in response to dsDNA damage. RPS27A and RPL40 are regulated post-transcriptionally and in a p53-independent manner, and their depletion persists days after the initial genomic lesion with Cas9. We also found that both non-specific double-strand breaks as well as single, targeted double-strand breaks reduce translation via eukaryotic initiation factor 2 alpha (eIF2α) phosphorylation, and that modulating the downstream effects of eIF2α phosphorylation during Cas9 editing leads to different repair outcomes. Ribosome profiling and RNA-seq data from Cas9-edited cells suggest that cells mount a translation response to dsDNA damage that precedes transcriptional changes. Our data demonstrate that Cas9-mediated genome editing can trigger temporary ribosome remodeling and translational shutdown in response to DNA double-strand breaks.

## Results

### Ribosome proteins RPS27A and RPL40 are downregulated after genome editing with Cas9

While investigating changes in ubiquitin gene expression after DNA damage, we serendipitously observed that the two ribosomal proteins encoded as fusion proteins with ubiquitin, RPS27A (eS31) and RPL40 (eL40), are downregulated after Cas9-guide RNA (gRNA) ribonucleoprotein (RNP) nucleofection (**Figure 1A**). This downregulation was apparent as late as 48-72 hours after nucleofection, even though at this point Cas9 was largely absent from the cell (**Figure 1B**) and genomic formation of indels was completed (**Figure 1C**). We found that RPS27A levels recovered 96 hours after nucleofection and RPL40 levels were beginning to increase within 72 hours (**Figure 1A**), suggesting that the cell resets protein expression three to four days after editing (**Figure S1A**).

**Figure 1.**
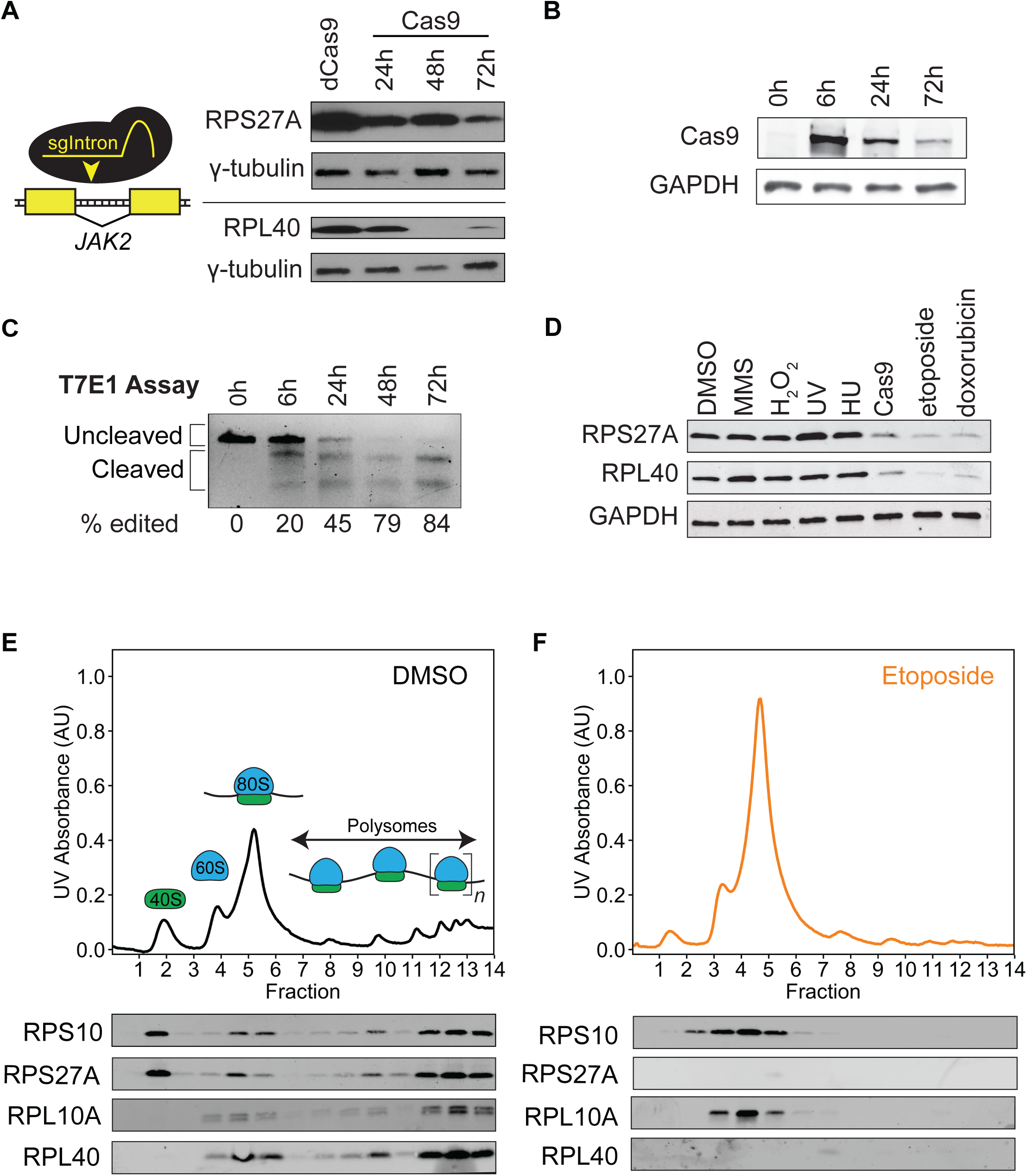
Ribosome proteins RPS27A and RPL40 are downregulated after genome editing with Cas9. A. Western blots reveal that RPS27A and RPL40 are depleted in HEK cells after nucleofection with Cas9 RNP complexes targeting intron 12 of *JAK2* (sgIntron). HEK cells harvested 72 hours post dCas9-sgIntron nucleofection served as the negative control.
B. Western blots depict rapid loss of Cas9 protein after RNP nucleofection.
C. T7 endonuclease 1 assay of JAK2 editing after Cas9-sgIntron nucleofection demonstrates that editing is largely complete after 24 hours. Band intensities were calculated using ImageJ, and percent edited was computed as 100% x (1-(1-fraction cleaved)^1/2^), where fraction cleaved = (sum of leavage product intensities)/(sum of uncleaved and cleaved product intensities).
D. Western blots of HEK cell lysates treated with different DNA damaging agents show that RPS27A and RPL40 are depleted after DNA double-strand breaks (DSB) and not other forms of DNA damage. MMS: methyl methanesulfonate, 0.03%, 1 hour. Cas9: Cas9*-*sgIntron nucleofection, 72 hr recovery. H_2_O_2_: hydrogen peroxide. UV: UV irradiation, 20 J/m^2^, 6 hour recovery. HU: hydroxyurea, 10 mM, 16 hours. Etoposide: 5 µM, 16 hours. Doxorubicin: 10 µM, 16 hours. DMSO: 0.01%, 16 hours.
E. Polysome profiles and ribosome protein Western blots of polysome profiling fractions from HEK cells treated with DMSO or (f) 5 µM etoposide for 16 hours reveal that RPL40 and RPS27A are lost from ribosome subunits after DSBs. UV absorbance = UV absorbance at 254 nm.

Downregulation of RPS27A and RPL40 depended on the DNA double-strand break, as catalytically inactive dCas9 did not provoke a similar response (**Figure 1A**). The guide RNA used in this experiment targeted a non-coding region of the *JAK2* gene (sgIntron), and *JAK2* levels remain unchanged after Cas9 nucleofection (**Figure S1B**). Our data therefore suggest that the loss of ribosomal subunits was due to the break itself and not disruption of *JAK2*. This days-long response was striking, as Cas9-mediated genome editing is often assumed to be relatively benign beyond the effects of the genomic sequence change itself.

We next asked whether ribosomal protein depletion was a specific response to DSBs versus other genomic lesions. We found that the loss of RPS27A and RPL40 does not occur after non-DSB DNA damage such as alkylation (methyl methanesulfonate), oxidative damage (hydrogen peroxide), thymine dimers (ultraviolet radiation), or replication fork stalling (hydroxyurea) (**Figure 1D**). By contrast, both single, targeted DSBs caused by Cas9 RNP nucleofection and multiple, unspecific DSBs induced by the topoisomerase II inhibitors etoposide or doxorubicin reduced RPS27A and RPL40 levels. Therefore, the loss of RPL40 and RPS27A we observed after genome editing is caused by multiple DSB-inducing agents and is specific to DSBs.

As RPS27A and RPL40 are core components of the ribosome, we wondered whether intact ribosomes lacked these core components or if the reduction in levels of these proteins reflected changes in the pool of free ribosomal subunits. We used Western blotting of polysome profiling fractions to measure the abundance of different ribosomal proteins in small (40S) and large (60S) ribosome subunits, 80S monosomes, and polysomes from cells treated with DMSO or etoposide (**Figure 1E-F**). Strikingly, etoposide caused an accumulation of 80S monosomes and a reduction of actively translating polysomes. We found that RPS27A and RPL40 were absent from 80S monosomes and other ribosomal subunits after etoposide treatment, while the control ribosomal proteins RPS10 (eS10) and RPL10A (uL1) remained. The lack of RPS27A and RPL40 in 80S monosomes and polysomes suggests that they are absent from actively translating ribosomes, but we cannot rule out the hypothesis that monosomes are not translationally competent after DSBs and that actively translating ribosomes require RPS27A and RPL40. In sum, we observe that DSBs cause an increase in 80S monosomes and reduction in translating polysomes, while RPS27A and RPL40 are lost from the translation machinery of etoposide-treated samples. Before investigating the DSB-induced accumulation of monosomes, we examined the mechanism by which RPS27A and RPL40 are lost after double-stranded DNA damage.

### Ubiquitins translated from *RPS27A* and *RPL40* decrease after dsDNA breaks

Since RPS27A and RPL40 are translated as polypeptide fusions between an N-terminal ubiquitin moiety and a C-terminal ribosomal protein, we asked if ubiquitin moieties associated with RPS27A and RPL40 are depleted from dsDNA-damaged cells. The ubiquitin-ribosomal protein fusions are post-translationally processed into separate polypeptides, and cleavage presumably occurs prior to incorporation of RPL40 and RPS27A into the ribosome, as the N-termini of RPL40 and RPS27A are positioned near the elongation factor binding site and the A site of the decoding center, respectively (Ben-Shem et al., 2010; Rabl et al., 2011). Ubiquitins translated from the four human ubiquitin genes, *RPL40* (also known as *UBA52*), *RPS27A*, *UBC*, and *UBB* are indistinguishable at the amino acid level, and consequently, we employed Cas9-mediated genome engineering to introduce unique epitope tags to the N-terminal ubiquitins associated with these loci. We created endogenously-tagged clonal cell lines for three of the four human ubiquitin genes: *V5-RPL40, HA-RPS27A*, and *Myc-UBC* (**Figure 2A**). Each of the ubiquitins encoded by these genes has an identical amino acid sequence, but the unique tag allows us to individually track them.

**Figure 2.**
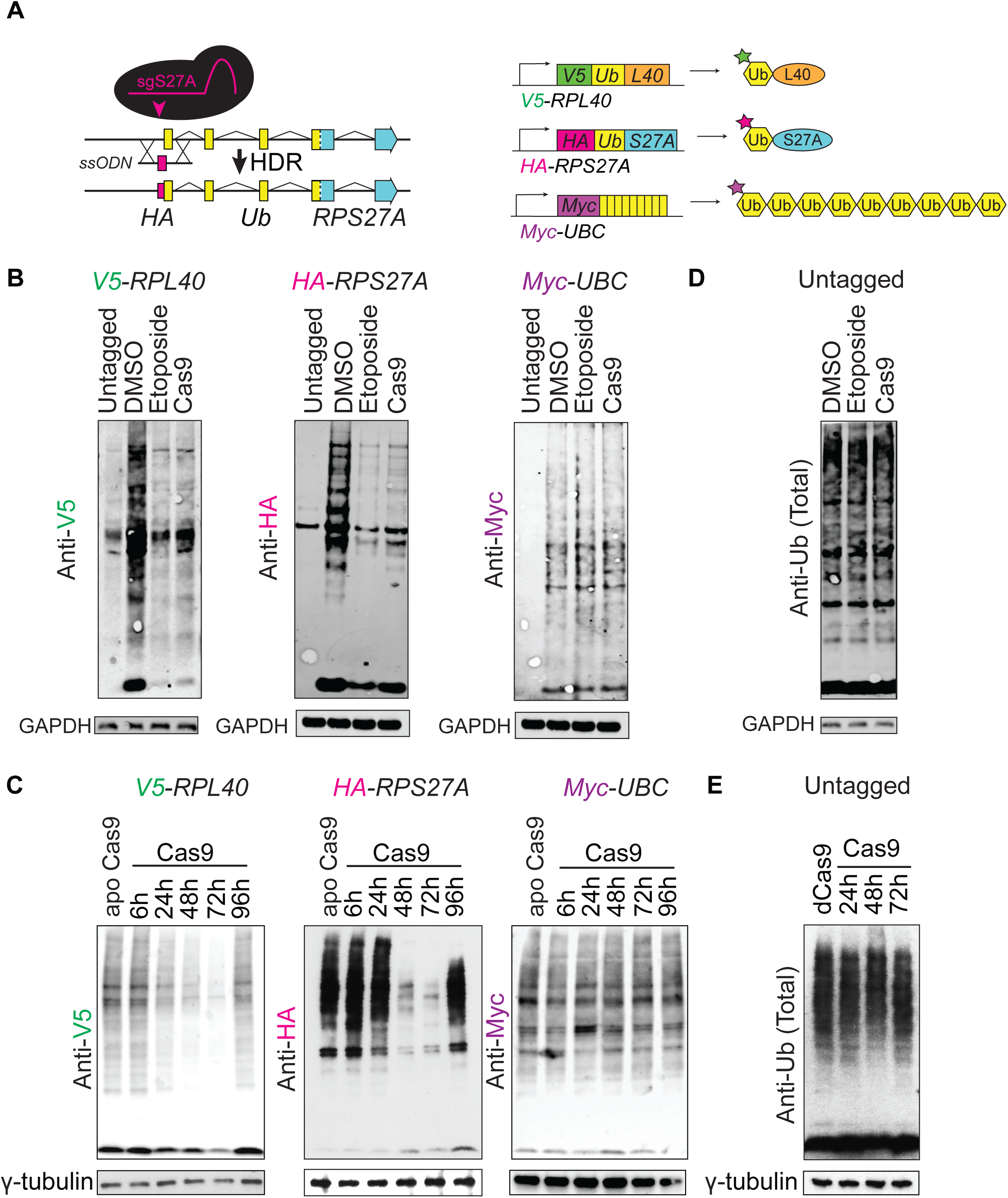
Ubiquitins translated from *RPS27A* and *RPL40* decrease after dsDNA breaks. A. Schematic of Cas9 genome editing strategy for introducing the HA epitope tag at the N-terminus of the ubiquitin translated from *RPS27A,* and a schematic of edited *HA-RPS27A, V5-RPL40,* and *Myc-UBC* and their primary translation products. *V5-RPL40* and *Myc-UBC* were edited in a similar fashion as *HA-RPS27A*. (ssODN = single-stranded oligodeoxyncleotide donor.)
B. Western blots depicting reduction of epitope-tagged ubiquitin translated from *V5-RPL40, HA-RPS27A*, but not *Myc-UBC,* 16 hours after treatment with 5 µM etoposide or 72 hours after Cas9-sgIntron nucleofection.
C. Western blots depicting the time course of depletion and recovery of epitope-tagged ubiquitin after Cas9-sgIntron nucleofection. apo Cas9 indicates Cas9 nucleofection without an sgRNA 72 hours post nucleofection.
D. Western blots show that total ubiquitin levels are unchanged after treatment with 5 µM etoposide or 72 hours after Cas9-sgIntron nucleofection.
E. Western blots show that there is no change in total ubiquitin levels 1-3 days after Cas9-sgIntron nucleofection. dCas9-sgIntron 72 hours after nucleofection served as the negative control.

Western blotting for each tag confirmed that the ubiquitin species from each edited locus could be uniquely tracked and incorporated into polyubiquitin chains (**Figure S2A**). We found that induction of either multiple DSBs with etoposide or a single DSB with a Cas9 RNP greatly reduced the abundance of the epitope-tagged ubiquitins translated from *V5-RPL40* and *HA-RPS27A* but had no effect on the ubiquitin associated with *Myc-UBC* (**Figure 2B**). By tracking tagged ubiquitin after a single Cas9 DSB, we found that the time course of *RPS27A* and *RPL40* ubiquitin depletion mirrored that of the RPS27A and RPL40 proteins, including recovery of the proteins several days after a DSB (**Figure 2C, Figure S2B**). Nucleofection of a targeted but catalytically inactive dCas9 RNP had no effect on the levels of the ubiquitins derived from *RPL40* or *RPS27A* (**Figure S2B**), confirming that the formation of a DSB was critical for loss of *RPS27A* and *RPL40*. Other forms of DNA damage such as MMS or UV radiation did not change the levels of ubiquitins associated with *RPL40* or *RPS27A* (**Figure S2C**). This mirrors the specificity to a double stranded DNA break we observed for the ribosomal proteins (**Figure 1D**), suggesting that the translation products of *RPL40* and *RPS27A* are repressed in tandem after dsDNA damage. Notably, DSBs had no gross effect on the total ubiquitin pool (**Figure 2E-F**), suggesting that cells are not modulating overall ubiquitin abundance.

### RPS27A is proteasomally degraded after dsDNA breaks

We next worked to identify the mechanism underlying the reduction in RPS27A and RPL40 after DSBs. We determined that loss of these proteins occurred post-transcriptionally, as qRT-PCR showed that DSBs induced by either etoposide or Cas9 did not affect the mRNA levels of *RPS27A* or *RPL40* (**Figure 3A-B**). In light of the key role played by ubiquitin signaling in proteasomal degradation, we wondered whether proteasomal degradation could explain the loss of RPS27A or RPL40. Indeed, we found that proteasome inhibition by epoxomicin treatment rescues the loss of RPS27A after DNA damage (**Figure 3C**). By contrast, the loss of RPL40 is unaffected, indicating that RPL40 is not proteasomally degraded after etoposide treatment (**Figure 3D**). Proteasome inhibition on its own increased basal RPS27A and RPL40 levels, suggesting some amount of constitutive degradation. The levels of other ribosomal proteins, including RPL22 and RPL10A, were unchanged by etoposide or epoxomicin treatment (**Figure 3E**). DSB-induced, proteasome-dependent degradation is therefore specific for RPS27A and does not globally affect the entire ribosome.

**Figure 3.**
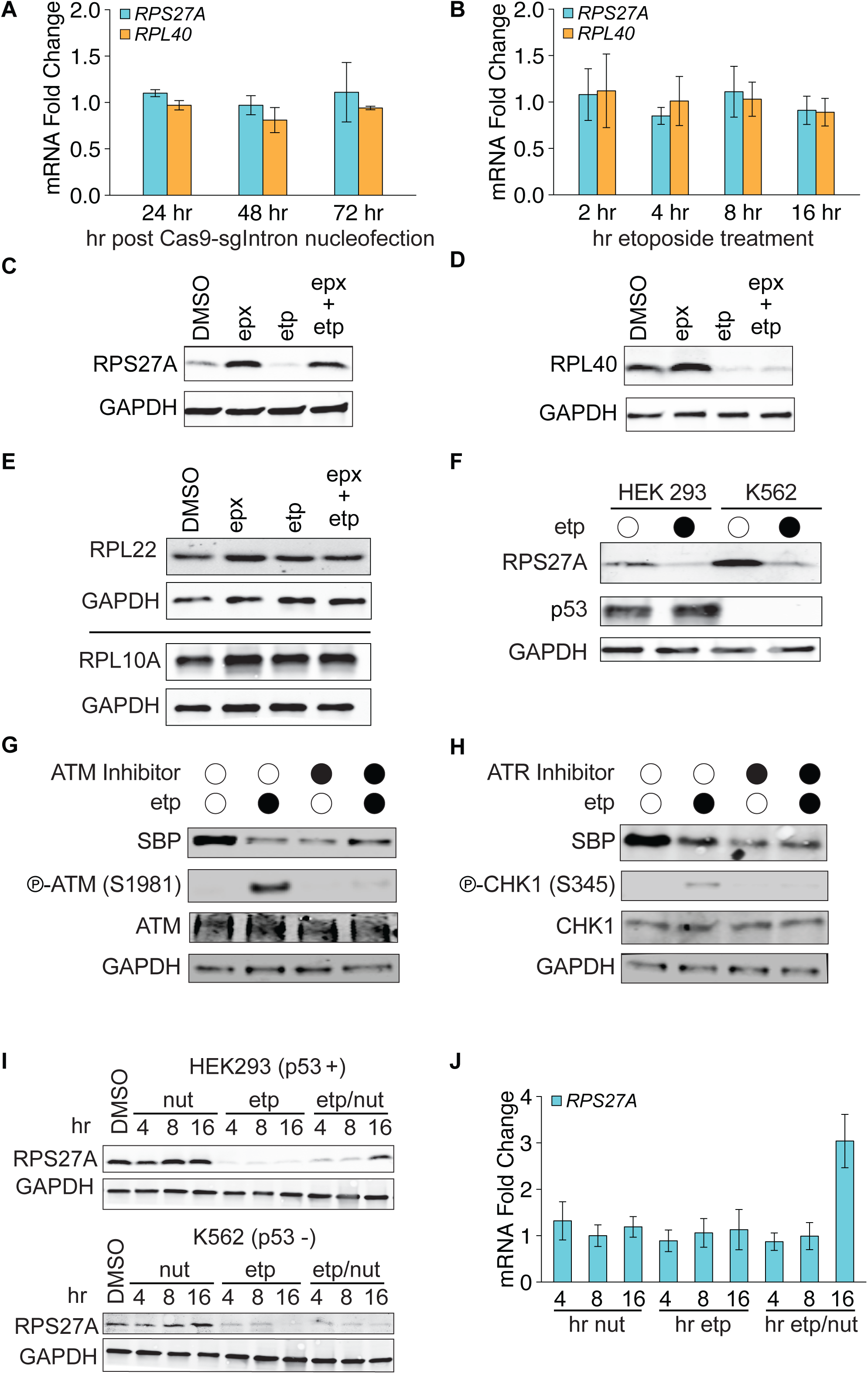
RPS27A is proteasomally degraded after dsDNA damage. A. Abundance of *RPS27A* and *RPL40* transcripts does not change after Cas9 RNP nucleofection. Fold changes were calculated using the 2^-ΔΔCt^ method with Cas9 without sgIntron (apo Cas9) as the control and *GAPDH* as the reference gene (n = 3, error bars = standard deviation).
B. As (A), showing abundance of *RPS27A* and *RPL40* transcripts does not change after 5 µM etoposide treatment for 16 hours.
C. Western blots demonstrate that proteasome inhibition with epoxomicin rescues RPS27A depletion.
D. As in (C), revealing that proteasome inhibition does not block DNA damage induced RPL40 depletion.
E. Western blots show that neither DNA damage nor proteasome inhibition affect levels of ribosomal proteins RPL22 or RPL10A.
F. Western blots confirm the p53 null status of K562 cells and demonstrate that loss of RPS27A is p53-independent.
G. ATM inhibition does not rescue RPS27A degradation. Transgenic RPS27A with a C-terminal SBP tag was followed by Western blotting after treatment with 5 µM etoposide and/or 10 µM ATM inhibitor, KU 55933. Phospho-ATM served as a positive control for ATM inhibition.
H. As in (G), showing that ATR inhibition (10 µM AZ 20 for 16 hours) does not rescue RPS27A degradation. Phospho-CHK1 served as a positive control for ATR inhibition.
I. Western blotting reveals a partial rescue of RPS27A levels in p53-positive HEK cells but not p53-null K562 cells when p53 degradation is inhibited with 10 µM nutlin.
J. *RPS27A* transcript abundance increases after DNA damage when p53 is stabilized by nutlin treatment (n = 4, error bars = SD).

Next, we wanted to test whether the proteasome-dependent loss of RPS27A reflected direct proteasomal degradation of RPS27A. We generated HEK Flp-In cell lines with single copy *Ub-RPS27A-SBP* or *RPS27A-SBP* transgenes lacking the endogenous promoter, introns, and UTR sequences. Both *RPS27A-SBP* and *Ub-RPS27A-SBP* generate protein products of the same molecular weight (**Figure S3A**), consistent with prior reports that the ubiquitin moiety is rapidly cleaved from RPS27A (Baker et al., 1992; Grou et al., 2015; Larsen et al., 1998). Since these transgenes are expressed in a non-native genomic context without most regulatory RNA elements, their loss after induction of a DSB further suggests post-transcriptional regulation.

We affinity purified RPS27A-SBP in denaturing conditions and used ubiquitin chain-specific antibodies to determine that RPS27A-SBP is basally modified with Lys48 polyubiquitin chains that signal for proteasomal degradation (Newton et al., 2008). Lys48 chain modification of RPS27A increases upon induction of DSBs with etoposide (**Figure S3B**). In contrast, we did not observe substantial modification of RPS27A-SBP by Lys63 or Met1 (linear) polyubiquitin chains, which generally do not target proteins to the proteasome. Taken together, our data indicate that cells lose mature RPS27A through proteasome-mediated degradation after dsDNA damage.

We next sought identify the DNA damage response pathway that triggers the degradation of RPS27A. Consistent with our observation that RPS27A is not regulated through transcription, we found that RPS27A degradation is independent of expression of p53; RPS27A is lost after DSBs in both p53-positive (HEK293) and p53-negative (K562) cell lines (**Figure 3F**). Using small molecule inhibitors, we found that the RPS27A response is not mediated through the activity of ATM or ATR, two of the master kinases that recognize damage at the site of the DSB and initiate a DNA damage response through a phosphorylation signaling cascade (Blackford and Jackson, 2017; Maréchal and Zou, 2013) (**Figure 3G-H**). Thus the upstream molecular signals that link DSB signaling with the depletion of RPS27A remain unclear.

Proteasomal degradation is initiated by ubiquitin ligases, which play a prominent role in several aspects of DNA damage signaling. MDM2 is a DNA damage regulated ubiquitin ligase that targets p53 for degradation under normal growth conditions and can also ubiquitinate RPS27A (Sun et al., 2011). However, we found that siRNA knockdown of MDM2 had no effect on the early loss of RPS27A caused by etoposide or Cas9-(**Figure S3C**). In contrast, we found that stabilizing p53 with the MDM2 inhibitor nutlin (Vassilev et al., 2004) rescued RPS27A levels at later time points after DSB formation, and this rescue was p53 dependent (**Figure 3I**). Recovery of RPS27A expression occurred at the transcriptional level (**Figure 3J**), consistent with *RPS27A* being a direct transcriptional target of p53 (Nosrati et al., 2015). We also found that nutlin rescued RPL40 levels after dsDNA damage and that this recovery is transcription dependent (**Figures S3D-E**). Overall, our data indicate that *depletion* of RPL40 and RPS27A is independent of p53 pathways, but the *reset* of levels of these proteins after DNA damage can be stimulated by p53.

We next turned towards a candidate approach to identify the ubiquitin ligase that regulates RPS27A. We first tested ZNF598, a mono-ubiquitin ligase known to ubiqutinate small ribosome subunit proteins RPS10 and RPS20 as part of the ribosome quality control pathway (Garzia et al., 2017; Sundaramoorthy et al., 2017). Knockdown of ZNF598 stabilized RPS27A in the presence of etoposide-induced DSBs, but had no effect on levels of RPL40 or other ribosome proteins, including the known ZFN598 target RPS10 (**Figure S3F**). In order to directly monitor RPS27A ubiquitination, we transiently expressed an epitope-tagged ubiquitin, immunoprecipitated this ubiquitin under denaturing conditions, and blotted for RPS27A. We observed ubiquitinated RPS27A under basal growth conditions, and its abundance increased upon proteasome inhibition and induction of DSBs with etoposide. Importantly, ZNF598 knockdown eliminated RPS27A ubiquitination, suggesting that ZNF598 is required for RPS27A ubiquitination (**Figure S3G**).

ZNF598 is a mono-ubiquitin ligase, but proteasomal degradation usually requires polyubiquitin Lys48 chains. As we previously found Lys48 polyubiquitin chains attached to RPS27A (**Figure S3B**), we postulated that another ubiquitin ligase extends the ZNF598-added monoubiquitin. This strategy of priming-and-extending by ubiquitin ligases has been previously described for other proteasomal substrates (Pierce et al., 2009; Wu et al., 2010). Given that MDM2 is not responsible for degradation of RPS27A, we tested the involvement of β-TRCP, which targets CReP, a eukaryotic initiation factor eIF2α phosphatase, for destruction after DNA damage (Loveless et al., 2015). We found that etoposide-induced RPS27A degradation is indeed rescued by knockdown of *β-TRCP* (**Figure S3H**). However, we found that depletion of ZNF598 or β-TRCP reduced etoposide-stimulated polyubiquitination of RPS27A to basal levels but did not eliminate ubiquitination (**Figure S3I**). Our data therefore cannot exclude regulation of RPS27A by ligases other than ZNF598 and β-TRCP. However, our data together with prior work on the molecular activities of ZNF598 and β-TRCP suggest a dual role for these ligases. We propose a ‘prime-and-extend’ model (Wu et al., 2010) in which RPS27A is first monoubiquitinated by ZNF598 and that this monoubiquitin is subsequently extended to Lys48-linked polyubiquitin chains by β-TRCP to signal proteasomal degradation of RPS27A.

### Double-strand DNA breaks lead to eIF2α phosphorylation and reduced translation initiation

Given the loss of RPS27A and RPL40 after dsDNA damage, we asked if cells exhibit a translation phenotype in response to DSBs. Consistent with our prior data (**Figure 1F**), polysome profiles of HEK293 cells treated with etoposide showed a sharp increase in 80S monosomes and a concordant reduction in polysomes (**Figure 4A**), demonstrating that etoposide-treated cells have fewer ribosomes per transcript. Etoposide-treated cells exhibited an imbalance in small (40S) and large (60S) ribosome subunits compared to DMSO-treated samples (40S:60S peak height ratio of 2:7 etoposide versus 1:1 DMSO), suggesting a deficiency in 40S subunits. Because the accumulation of monosomes is a hallmark of reduced protein synthesis, we wanted to gauge how nascent chain translation changes after DSBs. We used incorporation of L-azidohomoalanine (AHA), a methionine mimic that can be labeled with alkyne-conjugated probes, to track protein synthesis (Wang et al., 2017). Induction of multiple DSBs with etoposide led to a marked reduction in translation, consistent with accumulation of 80S monosomes (**Figure 4B**; **Figures S4A**). Surprisingly, induction of a single DSB with Cas9 led to reduced translation output as well (**Figure 4B**). Polysome profiling of Cas9 nucleofected cells revealed a modest increase in 80S, decrease in 40S, and shift from heavy to light polysomes (**Figure S4B**). Thus both chemically-induced DSBs and Cas9-mediated genome editing lead to a global reduction in protein synthesis.

**Figure 4.**
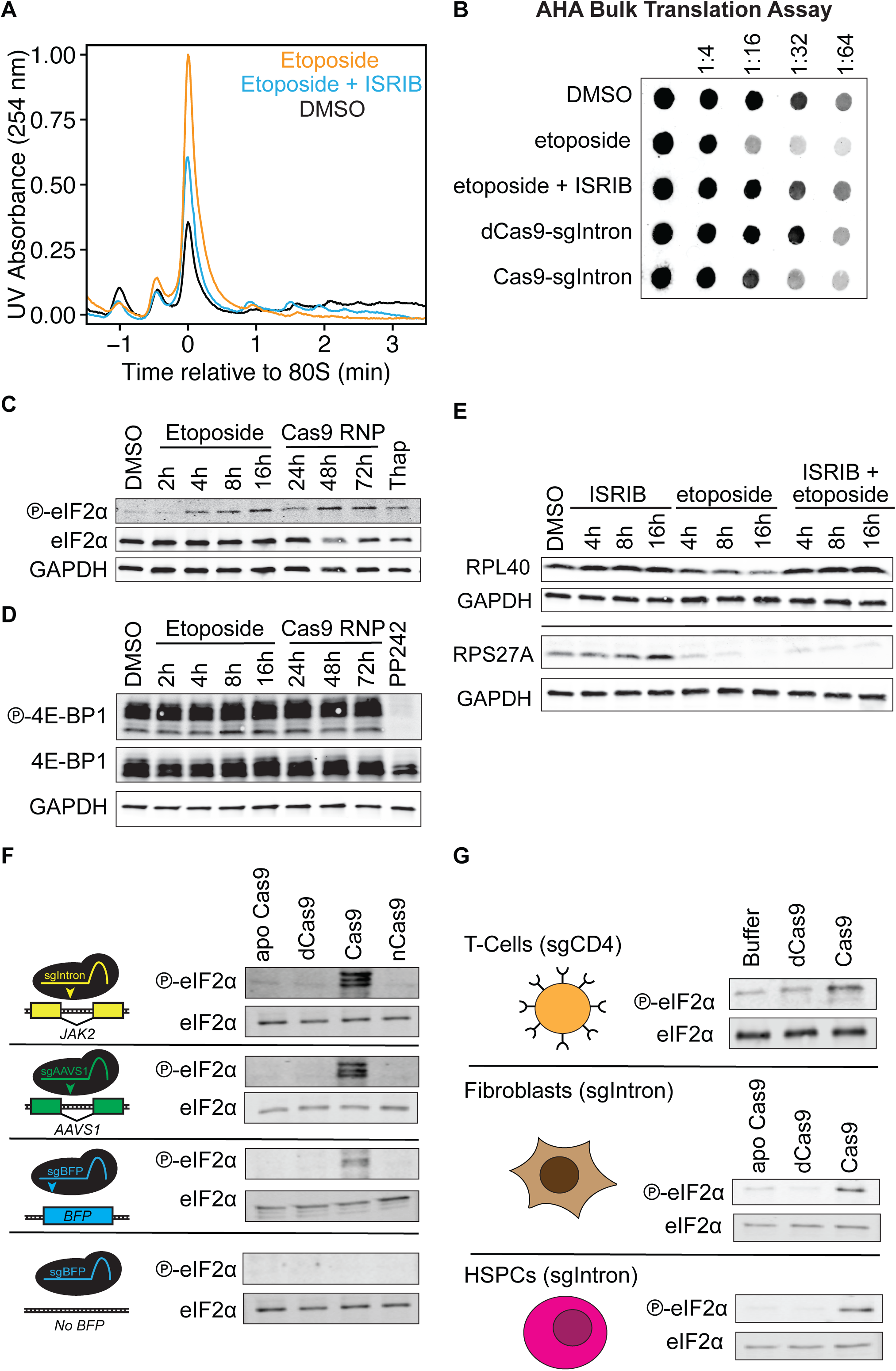
Double-strand DNA damaged leads to eIF2α phosphorylation and reduced translation initiation. A. Polysome profiles of HEK cells treated with 5 µM etoposide or 5 µM etoposide, 200 nM ISRIB for 16 hours.
B. AHA bulk translation assay demonstrates that dsDNA damage reduces protein synthesis. HEK cells were lysed after 16 hours after drug treatment or 72 hours after nucleofections with RNPs. Two hours before lysis, growth media replaced with methionine-free media containing a methionine mimic, L-azidohomoalanine (L-AHA). Lysates were normalized by protein content, labeled with IRDye 800CW-DBCO Infrared Dye, blotted on a nitrocellulose membrane, and imaged with a LI-COR Odyssey CLx Imager.
C. Levels of eIF2α (S51) phosphorylation increase in HEK cells treated with 5 µM etoposide or nucleofected with Cas9-sgRNA RNPs targeting a *JAK2* intron. Treatment with 1 µM thapsigargin (Thap) for 30 minutes served as a positive control for eIF2α phosphorylation.
D. As in (C), showing that levels of 4E-BP1 (T37/47) phosphorylation do not change after DNA damage. Treatment with 2.5 µM PP242 for 30 minutes served as a positive control for 4E-BP1 hypo-phosphorylation.
E. Western blotting indicates that phospho-eIF2α inhibitor ISRIB rescues loss of RPL40 but not RPS27A after DNA damage.
F. Only active Cas9 with a targeting gRNA triggers eIF2α phosphorylation. HEK or HEK-BFP cells were nucleofected with Cas9 without guide (apo Cas9), dCas9, Cas9, or nickase Cas9 and guides against a *JAK2* intron (sgIntron), the *AAVS1* locus (sgAAVS1), or a *BFP* transgene (sgBFP).
G. Western blotting reveals eIF2α phosphorylation in T-cells, fibroblasts, and MPB-CD34+ HSPCs 24 hours after nucleofection with sgCD4 or sgIntron RNPs.

We next asked if dsDNA-damaged cells regulate translation through either of two canonical mechanisms: the phosphorylation of eukaryotic initiation factor 2α (eIF2α) or the de-phosphorylation of 4E binding protein (4E-BP). Phosphorylation of eIF2α prevents eIF2 from recruiting the initiator methionine tRNA to the mRNA while de-phosphorylation of 4E-BP inhibits eIF4E from associating with the 5’ cap of transcripts (Jackson et al., 2010; Sonenberg and Hinnebusch, 2009). We found that multiple, non-specific etoposide-induced DSBs and a single, targeted Cas9-induced DSB both cause phosphorylation of eIF2α (**Figure 4C**). In contrast, we observed no changes in phosphorylation of 4E-BP (**Figure 4D**).

Phosphorylation of eIF2α translationally activates a group of transcripts collectively known as the integrated stress response (Sidrauski et al., 2013). We confirmed that etoposide increases expression of ATF4, a key integrated stress response transcription factor (**Figure S4C**). We also observed that co-administration with ISRIB, a small molecule that mitigates the downstream effects of eIF2α phosphorylation (Sidrauski et al., 2013), rescued the etoposide-induced accumulation of 80S monosomes, depletion of polysomes, and 40S:60S imbalance (**Figure 4A**), and restored bulk protein synthesis (**Figure 4B**). Our data indicate that both drug-and Cas9-induced dsDNA breaks lead to the inhibition of translation initiation through eIF2α phosphorylation.

We previously found that the etoposide-induced loss of RPL40 is not mediated through transcription or proteasomal degradation (**Figure 3A,D**), and we therefore asked whether RPL40 is regulated at the translational level by eIF2α signaling. We found that co-administration of ISRIB with etoposide completely prevented the loss of RPL40 caused by DSBs (**Figure 4E**). RPS27A levels were slightly increased by ISRIB in the presence of DSBs, but were far from completely rescued. Thus RPL40 is regulated at the translation level through a phosho-eIF2α dependent mechanism.

We next used Cas9 targeted to different genomic locations to explore whether eIF2α phosphorylation is a general response to genome editing. We tested guide RNAs that target the *JAK2* intron (sgIntron, see **Figure 1C** for editing efficiency), the *AAVS1* safe harbor site (sgAAVS1, (Richardson et al., 2016)) or a blue fluorescent protein (BFP) single-copy transgene (sgBFP, (Richardson et al., 2018)). All Cas9 RNPs caused eIF2α phosphorylation (**Figure 4F**). Nucleofecting Cas9 without a guide RNA (apo Cas9) had no effect on eIF2α phosphorylation, nor did nucleofection of guide RNAs complexed with catalytically inactive dCas9. Genomic nicking induced by the Cas9 D10A nickase (nCas9) also did not induce eIF2α phosphorylation. We confirmed that Cas9-induced eIF2α phosphorylation was specific to the dsDNA damage itself, as Cas9 RNP complexes only induced eIF2α phosphorylation when the guide RNA had a genomic target. When we nucleofected Cas9-sgBFP into parental HEK293 cells we found no evidence of eIF2α phosphorylation (**Figure 4F**), but nucleofecting the same RNP into HEK293 cells harboring a *BFP* transgene led to phosphorylation of eIF2α.

We also verified that eIF2α phosphorylation is a general response that occurs after Cas9 RNP editing in a range of primary cell types. Neither T-cells, hematopoietic stem and progenitor cells (HSPCs), nor fibroblasts exhibited high levels of eIF2α phosphorylation when nucleofected with negative control apo Cas9 or dCas9-sgRNA (**Figure 4G**; see **Figures S4D-E** for T-cell sgRNA target validation). However, nucleofection with catalytically active Cas9 complexed with multiple different targeting guide RNAs caused increased eIF2α phosphorylation in each of these primary cells. Primary cells are p53-positive, but we found that eIF2α phosphorylation also occurs in K562 p53-negative cells, much like RPS27A degradation (**Figure S4F**). Hence, a single locus Cas9-induced DSB triggers eIF2α phosphorylation in a wide range of cell types.

### Modulating eIF2α phosphorylation alters genome editing outcomes

Given that eIF2α phosphorylation is induced by DSBs, we wondered whether downstream eIF2α signaling influenced genome editing outcomes. We altered the eIF2α response using two small molecule drugs: ISRIB to bypass eIF2α signaling and salubrinal to increase eIF2α phosphorylation (**Figure S5A**, (Boyce et al., 2005)). We performed editing experiments with HEK293 or K562 cells treated with ISRIB or salburinal, targeting a single-copy BFP transgene in each cell line to introduce insertions and deletions (indels) via error-prone DNA repair. We monitored genome editing using both T7 endonuclease I (T7E1) heteroduplex assays and next-generation sequencing of PCR amplicons of the edited transgene.

Strikingly, increasing phospho-eIF2α signaling with salubrinal decreased the frequency of indels during Cas9-sgBFP editing. Bypassing phospho-eIF2α signaling with ISRIB, on the other hand, resulted in an increased fraction of indels (**Figure 5A-B**). Increasing eIF2α phosphorylation with salubrinal while simultaneously bypassing this phosphorylation with ISRIB overcame the salubrinal-induced decrease in editing (**Figure 5A-B**). Perturbing eIF2α signaling affected editing levels in both p53-positive (HEK) and p53-negative cells (K562) (**Figure 5A, S5B**). From next-generation sequencing of edited alleles, we found that modulating eIF2α phosphorylation changed the relative frequency of edited alleles rather than introducing new types of indels (**Figure 5C, Figure S5C, Table S1**). These data indicate that DSB-induced eIF2α signaling affects DNA repair to reduce the error-prone formation of indels.

**Figure 5.**
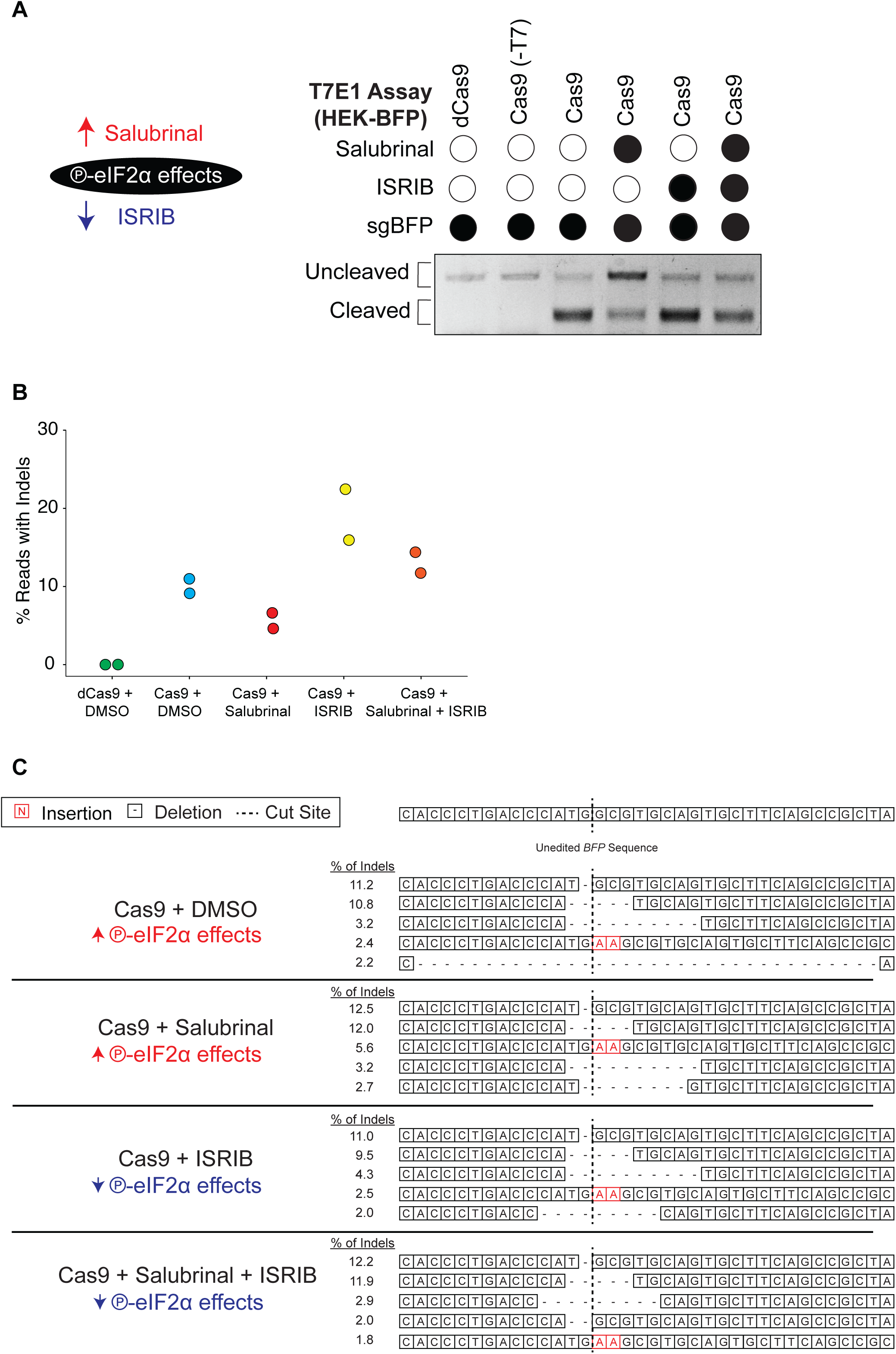
Modulating eIF2α phosphorylation alters genome editing outcomes. A. T7 Endonuclease 1 assay for genome editing of the transgenic *BFP* locus in HEK-BFP cells nucleofected with sgBFP-Cas9 (or dCas9) RNPs and treated with 75 µM salubrinal or 200 nM ISRIB for 16 hours. (Image analyzed as in **Figure 3A**).
B. Percentage of next generation sequencing (NGS) reads with insertions or deletions after genome editing, as in (A). Reads were aligned using NEEDLE (Li et al., 2015), and a modified version of CRISPResso (Pinello et al., 2016) was used to analyze editing outcomes.
C. Sequence identity and frequency of the top five *BFP* indel alleles from one of each experimental condition quantified in (B).

### Genome editing initiates a translational response that precedes long-term transcriptional changes

We wanted to measure how the ribosome remodeling and eIF2α phosphorylation induced by Cas9-mediated genome editing globally affect translation. We carried out ribosome profiling and matched mRNA sequencing in HEK293 cells with a single DSB induced by Cas9-sgIntron, with catalytically inactive dCas9-sgIntron serving as our background control (**Figure 6A**). *JAK2* mRNA and ribosome footprint levels did not show any significant differences at either 36 or 72 hours (**Table S2**), confirming our qPCR data (**Figure S1A**), which indicated that sgIntron-targeted editing does not perturb expression of *JAK2*. Global profiling of translation and transcription revealed that cells with a single Cas9-DSB activate an early translational program that is replaced by a longer-term transcriptional response. At 36 hours after nucleofection, we found 132 genes that exhibit changes in ribosome footprint abundance while no genes changed in transcript abundance (Wald test, FDR corrected *p*-value < 0.1, **Figures 6B-C**, **Table S2**). By 72 hours, there were changes in mRNA transcript levels but no statistically significant changes in footprint abundance (**Figures 6B&D**, **Table S2**). Translational efficiency, the ratio of ribosome footprints to mRNA transcripts, also reflected these differences, with changes in translational efficiency at 36 hours driven by translation and changes at 72 hours driven by mRNA abundance (**Figure S6A**).

**Figure 6.**
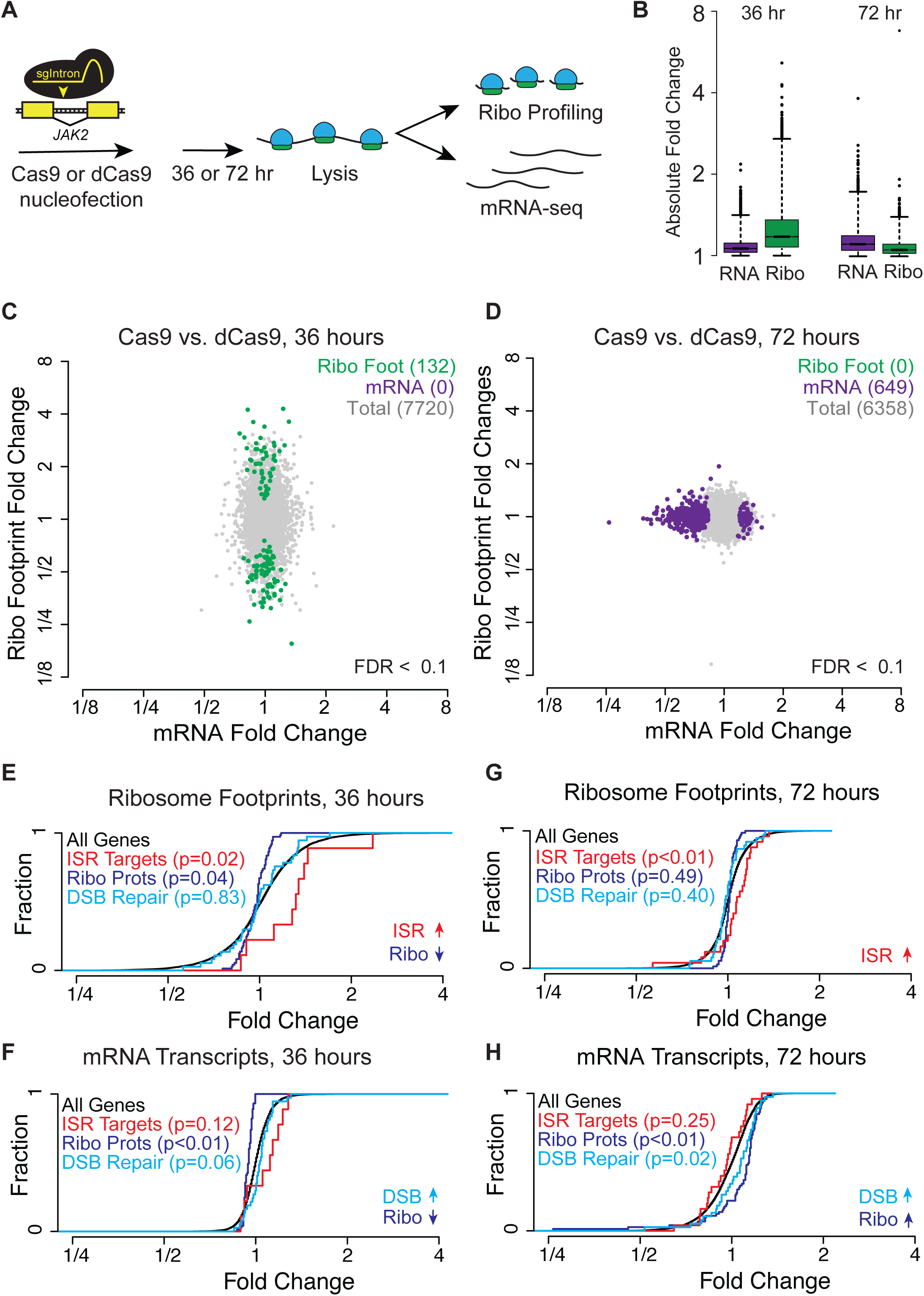
Genome editing initiates a translational response that proceeds long-term transcriptional changes. A. Experimental design for ribosome profiling and RNA-seq experiments. HEK cells were nucleofected with Cas9-sgIntron and harvested after 36 or 72 hours. Lysates were divided between ribosome profiling and RNA-seq experiments.
B. Distribution of absolute fold changes on a logarithmic scale, for genes identified in RNA-seq and ribosome profiling experiments at 36 and 72 hours post-editing. Whiskers denote values 1.5 * (the interquartile range).
C. Changes in ribosome footprint versus mRNA abundance (C) 36 hours or (D) 72 hours after Cas9 or dCas9 nucleofection. Green = genes with significant changes in ribosome footprints. Purple = genes with significant changes in mRNA transcripts (Wald test, FDR < 0.1).
D. Cumulative distribution function (CDF) plots for ribosomal protein genes (Ribo), integrated stress response targets (ISR), and DSB repair genes observed in the ribosome profiling (E-F) and mRNA-seq (G-H) experiments 36 hours (E and G) or 72 hours (F and H) after Cas9-sgIntron nucleofection. p-values were calculated using the Mann-Whitney-Wilcoxon rank sum test. See **Table S3** for target set gene lists.

Because we found that even a single DSB induces eIF2α phosphorylation, we asked whether genes known to be translationally regulated during the phospho-eIF2α-induced integrated stress response (ISR) also experience changes in translation after a Cas9-induced DSB. At both 36 and 72 hours after Cas9 nucleofection, we found that ISR targets (**Table S3**) collectively had higher translation (*p*< 0.05, Mann-Whitney-Wilcoxon test, **Figure 6E, 6G**), although individual genes did not rise to the level of significance. This effect was much larger at 36 hours than at 72 hours. Genome editing with Cas9 therefore leads to the induction of the integrated stress response at the translation level. These results provide a global view of cells activating translational and transcriptional responses that persist days after Cas9 is gone from the cell and genome editing is complete (**Figure 1B-C**).

Given that we observed changes in RPS27A and RPL40 levels after Cas9 editing, we asked how the global translation of ribosomal protein genes changes after a Cas9-mediated DSB. We found decreased footprints and mRNA abundance for several ribosomal protein transcripts 36 hours after Cas9 editing (*p* < 0.05, **Figure 6E-F**). eIF2α phosphorylation can lead to modest decreases in ribosome protein translation (Sidrauski et al., 2015), and our data links this eIF2α signaling to the DSB response. Ribosome protein transcript levels increased 72 hours after a Cas9-mediated DSB, suggesting that the cell resets ribosome protein levels through increased transcription (**Figures 6G-H**). Given our previous data that the reset of *RPS27A* and *RPL40* transcripts after DSBs is p53-dependent (**Figure 3I, S3D**), it is tempting to speculate that the global transcriptional increase in ribosomal transcription is the result of p53 signaling.

In the Cas9 ribosome profiling datasets, we found that DSB repair genes are somewhat regulated at the translation level. DSB repair genes showed no significant change in translation at 36 hours (**Figures 6E, S6B**) but showed a small decrease in translation efficiency at 72 hours that was driven by transcript abundance (**Figures S6B**). This decrease in translation may signify that the cell tunes down the production of these proteins as the cell returns to homeostasis. Our data, however, do not exclude early translational control of DSB repair genes that is completed before 36 hours. In sum, our ribosome profiling and RNA-seq data from Cas9-treated cells demonstrate that even a single DSB can induce small, yet significant changes to the translatome and transcriptome that persist days after the lesion is formed and repaired. Overall, our data suggest that Cas9 editing leads to changes in signaling, translation, and gene expression that are not only independent of editing a target gene but also inherent to the cellular response to double stranded DNA damage.

## Discussion

DNA damage poses a serious threat to organisms. Consequently, cells have an array of damage response pathways dedicated to maintaining genome integrity. These responses include cell cycle arrest after moderate levels of damage and apoptosis when the insult becomes too great. One hallmark of the DNA damage response is transcriptional reprogramming, such as the p53 response. Here, we report another, translational layer of DSB response. Even a single DSB caused by Cas9 genome editing can induce potent, p53-independent ribosome remodeling and translational reprogramming that occurs prior to transcriptional changes.

### Translational shutdown after DNA damage promotes error-free repair

We found that DSBs introduced during genome editing lead to translational reprogramming in immortalized and primary human cell types. Bulk protein synthesis is reduced after DSBs in part because translation is inhibited by eIF2α phosphorylation. Other types of DNA damage can induce eIF2α phosphorylation (Deng et al., 2002; von Holzen et al., 2007; Jiang and Wek, 2005; Kim et al., 2014; Peidis et al., 2011; Robert et al., 2009; Wu et al., 2002), and we found that multiple DSBs or even a single DSB leads to eIF2α phosphorylation. However, single-strand genomic lesions do not induce this signal (Cas9 vs. nickase Cas9,**Figure 4D**). Ionizing radiation can cause mTOR-mediated dephosphorylation of 4E-BP (Braunstein et al., 2009; Kumar et al., 2000; Schneider et al., 2005), but we found no evidence that cells with chemically-or Cas9-induced DSBs reduce translation through 4E-BP dephosphorylation (**Figure 4C**). This difference may reflect other cellular responses to the collateral damage caused by ionizing radiation to non-DSB DNA lesions or to other macromolecules including RNA and protein. We have found that the DSB translational response does not require canonical DNA damage factors such as p53. Reset of ribosomal protein levels after DSBs can be stimulated by p53-mediated transcription (**Figure 3J, S3E**), but the upstream signaling pathways linking DNA damage to eIF2α phosphorylation remain unclear.

We found that eIF2α phosphorylation may help cells avoid permanent genomic changes after double stranded DNA damage. Notably, bypassing eIF2α phosphorylation increases error-prone repair at a Cas9 DSB, while increasing eIF2α phosphorylation decreases indel formation. Cells have a powerful incentive to avoid error-prone repair, and it has been suggested that nonhomologous end joining (NHEJ) is inherently a fidelitous process (Boulton and Jackson, 1996; Honma et al., 2007; Lin et al., 2013; Rath et al., 2014). In this model, the indels caused by genome editing are products of non-fidelitous alternative end joining (alt-EJ) pathways such as microhomology mediated end joining (MMEJ) or processing of the DNA ends prior to repair (Bae et al., 2014; Bétermier et al., 2014; Guirouilh-Barbat et al., 2007; Nakade et al., 2014). It is tempting to speculate that DSB-induced eIF2α phosphorylation could promote error-free DNA repair as a means to maintain genome fidelity. However, the downstream players that alter the repair profile of a genomic locus after eIF2α phosphorylation remain to be identified.

### Translational changes bridge the immediate, post-translational DNA damage response to the long-term transcriptional response

Double stranded DNA breaks elicit an immediate post-translational response that enacts immediate processing of the break. This response includes phosphorylation of proteins such as ATM and H2AX and ubiquitination of proteins such as p53 and histones. DSBs also induce a potent p53-mediated transcriptional response, leading to reprogramming that prioritizes DNA damage response. We have found that DSBs induce a short-term translational response mediated by eIF2α phosphorylation.

We hypothesize that the translational response to DNA damage enables cells to bridge the immediate post-translational response with longer-term transcriptional reprogramming. Cells increase the translation of integrated stress response genes 36 hours after Cas9 RNP nucleofection, suggesting that cells activate a translational program to cope with DNA damage prior to transcriptional changes. We find that this translational program is shut off by 72 hours, with changes in mRNA levels dominating gene expression.

We observed that RPS27A and RPL40 could be stimulated by p53-mediated transcription after DSBs (**Figures 3J, S3E**), consistent with reports that *RPS27A* can be a transcriptional target of p53 (Nosrati et al., 2015). RPS27A was previously described as binding and inhibiting the E3 ligase MDM2 (Sun et al., 2011), thereby promoting p53 expression in the cell. The role of RPS27A in preventing p53 degradation coupled with p53 activation of *RPS27A* transcription suggests an RPS27A-p53 positive feedback loop. Consequently, the degradation of RPS27A may serve to keep this loop inactive or shut it off after repair.

### Ribosomes lack core ribosome proteins after dsDNA damage

Non-DSB DNA damage caused by sources such as UV irradiation and cisplatin leads to the inhibition of Pol I transcription (Ciccia et al., 2014; Kruhlak et al., 2007; Larsen et al., 2014), preventing rRNA expression and impacting ribosome biogenesis. Interestingly, Cas9 or I-PpoI-induced DSBs in rDNA triggers this inhibition (van Sluis and McStay, 2015). Our study has revealed that DSBs lead to translation phenotypes beyond impaired ribosome biogenesis regardless of their location in the genome. Our observation that ribosomes lack RPL40 and RPS27A after dsDNA damage is one of the few known instances where ribosome composition is deliberately modulated in response to a specific biological stimulus (Shi and Barna, 2015; Xue and Barna, 2012). While differential expression of ribosomal proteins between tissue types and subpopulations of ribosomes within a cell are emerging themes in ribosome biology, there have been few cases of altered ribosome composition in response to the cellular environment.

While loss of RPS27A and RPL40 may alter ribosome function in a way that is difficult to detect in our ribosome profiling analysis, we cannot rule out that ribosomes lacking RPS27A and RPL40 have different functions. Indeed, ribosomes lacking RPL40 are capable of translation in certain contexts. RPL40 is necessary for vesicular stomatitis virus (VSV) translation but not cap-dependent translation in HeLa cells (Lee et al., 2012). In fact, complete deletion of the paralogous *RPL40A* and *RPL40B* genes in yeast was not lethal, and affected translation of only ∼7% of the genome. RPL40 depletion – and perhaps RPS27A depletion as well – may thus act in a regulatory fashion. It is also possible that changes in ribosome composition after DNA damage may serve at least in part to regulate the extra-translational functions of RPS27A, RPL40, or their associated ubiquitins.

### Gene editing induces cellular phenotypes

There is growing appreciation that Cas9 genome editing can cause cellular effects that mirror those observed with multiple, non-specific DSBs. The degree of damage may be far less, but the principle is the same. For example, embryonic stem cells are hyper-sensitive to HDR from even a single DSB introduced by Cas9, which can induce a p53 response that compromises cell health (Haapaniemi et al., 2018; Ihry et al., 2018). CRISPR-Cas9 nuclease screening data has also shown that targeting high copy number or repetitive regions of a genome reduces cell fitness, consistent with a titratable cell cycle arrest that could be caused by p53 signaling (Aguirre et al., 2016; Munoz et al., 2016; Wang et al., 2015).

We have found that even a single non-coding Cas9-induced DSB elicits ribosome remodeling and translational shutdown. Much of the concern about the safety and efficacy of genome editing had focused on off-target mutagenesis. Our findings highlight how the endogenous DNA damage response can have a days-long impact on the translatome and transcriptome independent of the gene target. These cellular responses should be taken into account when it is impossible to isolate and expand a clonal cell line for long periods of time after genome editing, for example during therapeutic genome editing of primary cells.

## Supporting information

## Acknowledgements

We thank Gloria Brar and Jamie Cate for their helpful discussions, and we also thank the Rape Lab at UC Berkeley for providing us with the pHA-Ub plasmid and the Cate Lab for supplying us with the thermophilic yeast strain, *Kluyveromyces marxianus.* We would like to acknowledge the QB3 MacroLab at UC Berkeley for purifying Cas9 proteins and the *Kluyveromyces marxianus* deadenylase; the UC Berkeley DNA Sequencing Facility for Sanger sequencing analysis; and the Vincent J. Coates Genomics Sequencing Laboratory for performing our NGS sequencing.

## Author Contributions

Conceptualization, JEC, NTI, EZ, and CR; Methodology, JEC, NTI, EZ, CR, SKW, AM, DNN, and JRL; Formal Analysis, NTI, SKW, CR, and EZ; Investigation, EZ, CR, DNN, PAF, JRL, ZAM, and JTV; Writing - Original Draft, CR and JEC; Writing - Review & Editing, CR, JEC, NTI, EZ, DNN, AM, and PAF; Visualization, CR, EZ, SKW, NTI, and DNN; Supervision, JEC, NTI, AM, and CR; Funding Acquisition, JEC, NTI, CR, AM, and DNN.

## STAR Methods

### CONTACT FOR REAGENT AND RESOURCE SHARING

Further information and requests for resources and reagents should be directed to and will be fulfilled by the Lead Contact, Jacob Corn.

### EXPERIMENTAL MODEL AND SUBJECT DETAILS

#### Cell Culture - Immortalized Cell Lines

HEK 293 (ATCC) and HEK Flp-In T-Rex cell lines (Invitrogen) were cultured in DMEM, high glucose, GlutaMAX (Gibco) with 10% FBS (VWR) in a 37°C incubator with 5.0% CO_2_ and 20% O_2_. K562 cells (ATCC) were cultured in RPMI (Gibco) with 10% FBS (VWR), 10% sodium pyruvate. Human neonatal dermal fibroblasts (ScienCell, Cat# 2310) were cultured in DMEM, high glucose with 10% FBS, 0.01% BME, 1% NEAA, 1% Sodium Pyruvate, 1% Glutamax, 1% HEPES, 1% pen/strep. Mobilized Peripheral Blood CD34+ Stem/Progenitor Cells (AllCells) were cultured in StemSpan^™^ Serum Free Expansion Media II (STEMCELL Technologies), StemSpan^™^ StemSpan(tm) CC110 (STEMCELL Technologies), 1% Pen/Strep.

#### Primary T-cell Isolation and Stimulation

Primary human T cells were isolated from two de-identified healthy human donors from Trima Apheresis leukoreduction chamber residuals (Vitalant, formally Blood Centers of the Pacific). Peripheral blood mononuclear cells (PBMCs) were isolated by Ficoll centrifugation using SepMate tubes (STEMCELL, per manufacturer’s instructions) then stored frozen in BAMBANKER serum-free freezing medium (Lymphotec Inc, per manufacturer’s instructions) until use. PBMCs were thawed and CD4+ T cells were further isolated by magnetic negative selection using an EasySep Human CD4+ Cell Isolation Kit (STEMCELL, per manufacturer’s instructions). Immediately following isolation, CD4+ T cells were then stimulated for 2 days by culture at initial concentration 1 x 10^6^ cells/mL in XVivo15 medium (STEMCELL) with 5% Fetal Bovine Serum, 50 mM 2-mercaptoethanol, and 10 mM N-Acetyl L-Cystine together with anti-human CD3/CD28 magnetic Dynabeads (ThermoFisher) at a beads to cells ratio of 1:1, along with a cytokine cocktail of IL-2 at 200 U/mL (UCSF Pharmacy), IL-7 at 5 ng/mL (ThermoFisher), and IL-15 at 5 ng/mL (Life Tech).

## METHOD DETAILS

### Cas9 RNP Nucleofection

gRNAs were in vitro transcribed as previously described (DeWitt et al., 2016; Lingeman et al., 2017). In brief, gRNA transcription template contain a T7 RNA pol promoter followed by target specific region and constant region (T7FwdVar) along with primer that is reverse complement of the invariant region of T7FwdVar (T7RevLong) and amplification primers (T7FwdAmp and T7RevAmp). Transcription templates for gRNA synthesis were PCR amplified from the primer mix. Phusion high fidelity DNA polymerase was used for assembly (New England Biolabs). Assembled template was used without purification for in vitro transcription by T7 polymerase using the HiScribe T7 High Yield RNA Synthesis Kit (New England Biolabs). RNA was purified with RNeasy kit (Qiagen). Cas9, dCas9, and D10A Cas9 (nCas9) proteins were purified using the protocol detailed in (Lingeman et al., 2017). Cas9, dCas9, and D10A Cas9 ribonucleoproteins (RNPs) were prepared as detailed in (Lingeman et al., 2017) with the exception of the T-cells experiments (see “Cas9 RNP Nucleofections with T-cells”). IVT gRNAs were used in all experiments except for the 36-hour ribosome profiling experiment, which used synthetic gRNA (Synthego), and the T-cell nucleofections, which used synthetic crRNAs and tracrRNAs (Dharmacon).

HEK cells were passaged 2 days before nucleofection and trypsinized at 60-90% confluency. For RNP nucleofections, either 100 pmol Cas9 and 120 pmol gRNA were added to 2.5 x 10^5^ cells in 20 µl SF Solution (Lonza), or 300 pmol Cas9 and 300 pmol guideRNA were added to 1 x 10^6^ cells suspended in 100 µl SF Solution (Lonza). HEK cells were nucleofected using program CM-130 in the X Unit of a Lonza 4D-Nucleofector (AAF-1002X, AAF-1002B) and pre-warmed media was immediately added to the cuvettes to increase cell viability. K562 cells were nucleofected with Cas9 RNPs as described for HEK cells using buffer SF and program FF-120; fibroblasts were nucleofected using buffer P3 and program DT-130. For HSPC nucleofections, 3,000 cells were nucleofected 30 pmol Cas9 and 36 pmol gRNA in solution P3 using pulse code ER-100 and recovered in 96-well plate.

### Cas9 RNP Nucleofections with T-Cells

RNPs were produced by complexing a two-component gRNA to Cas9. A crRNA targeting exon 2 of the human *CD4* gene (UUGCUUCUGGUGCUGCAACU, (Hultquist et al., 2016)) and tracrRNA were chemically synthesized (Edit-R, Dharmacon). Lyophilized RNA was resuspended in 10 mM Tris-HCL (7.4 pH) with 150 mM KCl at a concentration of 160 µM, and stored in aliquots at −80 °C. Recombinant Cas9-NLS or dCas9-NLS were purified as detailed in (Lingeman et al., 2017) and stored at 40 µM in 20 mM HEPES-KOH pH 7.5, 150 mM KCl, 10% glycerol, 1 mM DTT. crRNA and tracrRNA aliquots were thawed, mixed 1:1 by volume, and annealed by incubation at 37°C for 30 min to form an 80 µM gRNA solution. Cas9 or dCas9 was then mixed with freshly-annealed gRNA at a 1:1 volume ratio (2:1 gRNA to Cas9 molar ratio) then incubated at 37 °C for 15 min to form a ribonucleoprotein (RNP) at 20 µM. RNPs were electroporated immediately after complexing.

Stimulated CD4+ T cells were harvested from their culture vessels and magnetic anti-CD3/anti-CD28 Dynabeads were removed by placing cells on an EasySep cell separation magnet (STEMCELL) for 4 minutes. Immediately prior to electroporation, cells were centrifuged for 10 minutes at 90 x g, then resuspended in the Lonza electroporation buffer P3 at a concentration of 5.0 x 10^7^ cells per mL. One million CD4+ T cells (20 µL) and 3 µL of Cas9-NLS RNP, dCas9-NLS RNP, or Tris buffer were added to each well of a 96-well electroporation plate (Lonza) in three replicates for each condition for each of two cell donors. Electroporation was performed with a Lonza 4D 96-well electroporation system with pulse code EH115. 15 minutes following electroporation, each well was split between three replicate 96-well plates and cultured in XVivo15-base growth medium (as above) supplemented with 500 U/mL IL-2 at an approximate density of 1 x 10^6^ cells per mL of media.

Approximately 20 hours after electroporation, lysates were prepared for Western blot analysis from samples from one replicate plate of edited T-cells. Cells were collected and centrifuged at 300 x g for 5 minutes. Culture media was aspirated off the cells, and cells were resuspended in PBS. This was repeated for a total of 2 PBS washes. After the final wash, cells were resuspended in 1X RIPA Lysis buffer (Cell Signaling Technologies) with Protease Inhibitor and Phosphatase inhibitor (Cell Signaling Technologies), incubated 10 minutes on ice, then stored at −80 °C.

Three days after electroporation, samples from a second replicate of edited T cells was collected and stained with Anti-CD3-PE (clone UCHT1, Biolegend), Anti-CD4-PECy7 (clone OKT4, Biolegend), and GhostDye780 (Tonbo). Fluorescence was measured on an Attune NxT Flow Cytometer (Thermo Fisher Scientific) and analyzed using FlowJo (Treestar, Inc) for presence or knockdown of surface expression of CD4.

### Western Blotting

Cells were pelleted at 400 x g for 5 minutes then washed twice with PBS before being lysed in RIPA buffer (1% SDS, 50 mM Tris HCl, pH 8.0 with 1X Halt Protease Inhibitor Cocktail or 1X Halt Protease and Phosphatase Inhibitor Cocktail, (Thermo Scientific). Lysates were incubated for 30 minutes on ice, vortexed for 30 seconds, and spun at 18,000 x g for 10 minutes at 4°C. Lysates were normalized using either BCA Assay (Pierce) or a Bradford Assay (Proteomics Grade, VWR) before being boiled at 97°C for 5 minutes with Laemmli buffer or Novex LDS Sample Buffer (Thermo Fisher Scientific). Samples were loaded onto NuPAGE 4-12% Bis-Tris Gels (Invitrogen) or Mini-Protean TGX precast gels (Bio-Rad) and run for 200V for 40 minutes.

Proteins were transferred onto nitrocellulose membranes using the Trans-Blot Turbo Blotting System (Bio-Rad) according to the manufacturer’s protocol. Membranes were blocked in 5% milk in TBST, washed 3 x 5 minutes in TBST, and incubated with primary antibodies in TBST with 5% BSA overnight at 4°C. Membranes were washed 3 x 5 minutes in TBST and incubated with either IRDye 800CW (LI-COR), IRDye 680RD (LI-COR), or HRP-conjugated secondary antibodies in 5% milk for 40 minutes before 2 x 5 minute washes with TBST and 1 x 5 minute wash with PBS. Blots were imaged by a LI-COR Odyssey CLx Imager or Pierce ECL reagents (Thermo Fisher) and X-ray film. All primary antibodies were used at a 1:1000 dilution except for anti-p53 (1:500, Santa Cruz Biotechnology, Cat# sc-126) and anti-phospho-ATM (1:50,000, Abcam, Cat# ab81292). See Key Resources Table for the complete list of antibodies.

### T7 Endonuclease 1 Assay

Edited cells were gathered off of plates with a pipette, spun at 10,000 x g for 1 min, washed once with PBS, and lysed in QuickExtract^™^ DNA Extraction Solution (Lucigen). Lysates were incubated at 65°C for 6 minutes and 98°C for 2 minutes in a thermocycler. Edited regions were PCR amplified in 100 ul reactions with AmpliTaq Gold 360 Master Mix (Thermo Fisher Scientific). PCR products were purified using MinElute PCR Purification Kit (Qiagen). PCR products were hybridized and digested with T7 endonuclease 1 (NEB) according to the NEB protocol for determining genome targeting efficiency. Digests were run on a 2% agarose gel. Relative intensities from DNA bands were quantified using ImageJ (Schindelin et al., 2015) with % edited = 100 x (1-(1-fraction cleaved)^1/2^) where fraction cleaved = (sum of cleavage product intensities)/(sum of uncleaved and cleaved product intensities).

### Inducing DNA Damage

For chemically inducing double-strand DNA damage, HEK cells were grown to 70% confluency then treated for 16 hours with 5 µM etoposide (Sigma-Aldrich) or 10 µM doxorubicin (Sigma-Aldrich). For chemically inducing other forms of DNA damage, HEK cells were treated with 0.03% methyl methanesulfonate (MMS) for 1 hour, 500 µM hydrogen peroxide for 1 hour, or 10 mM hydroxyurea for 16 hours. To damage cells using ultraviolet light, cells were irradiated at 20 J/m^2^ with a FB-UVXL-1000 UV Crosslinker (Fisher Scientific) and recovered for 6 or 24 hours before lysis. Cells were treated with DMSO for 16 hours as a negative control unless otherwise noted.

### Polysome Profiling

HEK cells cultured in 10 cm plates were washed with 10 ml DPBS before lysis with ice cold 100-400 µl polysome buffer (20 mM Tris pH 7.4, 150 mM NaCl, 5 mM MgCl_2_, 1 mM DTT, 100 µg/mL cycloheximide) with 1% Triton X-100 and 25 U/ml TURBO DNase (Thermo Fisher Scientific). When polysome profiling fractions were collected for protein analysis, 2X protease inhibitor cocktail (P1860, Sigma) was added to the lysis buffer and sucrose gradients. The amount of cells varied between experiments but was generally between 1-8 x 10^6^ cells per biological condition. Cells were scraped off plates in lysis buffer and incubated on ice in microcentrifuge tubes for 10 minutes. Lysates were spun at 10 minutes at 20,000xg, and the supernatants were normalized using the Quant-iT RiboGreen RNA Assay Kit (Thermo Fisher Scientific).

6 ml 50% (w/v) sucrose in polysome buffer was layered under 6 ml 10% sucrose solution in 14 x 89 mm ultracentrifuge tubes (VWR), and 10-50% sucrose gradients were created using a Gradient Master (BioComp Instruments) with rotation set at 81.5°, speed 16 for 1:58. 200 µl normalized cell lysate (with RNA concentrations generally between 50-250 ng/µl) was layered on top of the gradients, and the gradients were loaded into Beckman Sw41 Ti rotor buckets and spun at 36,000 rpm (∼250,000xg) for 2.5 or 3 hours at 4°C in a Beckman L8-M Ultracentrifuge. Sucrose gradients were pumped through the Gradient Master at 0.2 mm/s, and UV absorbance at 254 nm was measured using a BioRad EM-1 Econo UV Monitor connected to a laptop running the Logger Lite software package (Vernier). Depending on the downstream experiment, fractions were manually collected every 20 to 24 seconds for a total of 15-18 fractions per sucrose gradient. Proteins were extracted for Western blots using methanol/chloroform extraction as detailed in Click-it Metabolic Labeling Reagents for Proteins (Invitrogen), and pellets were boiled at 95°C in 1X Laemmli buffer before SDS-PAGE.

### RT-qPCR

RNA was extracted from cells using the Direct-zol^™^ RNA MiniPrep Kit (Zymo) according the manufacturer’s instructions. 1 µg total RNA was used for reverse transcription with Superscript III First Strand Synthesis SuperMix (Thermo Fisher Scientific). qRT-PCR was performed using Fast SYBR Green Master Mix (Applied Biosystems) on a StepOnePlus Real-Time PCR System (Applied Biosystems). C_t_ values from target genes were normalized to *GAPDH*, and the expression of each gene was represented as 2^-(ΔΔCt)^ relative to the reference sample.

### Endogenous Tagging of Ubiquitin Genes

We used Cas9 genome editing to endogenously tag the N-terminal ubiquitins of *RPS27A, RPL40* (also known as *UBA52*), and *UBC* genes with the HA, V5, and Myc tags, respectively. We designed gRNA sequences upstream of the *RPS27A, RPL40,* and *UBC* ubiquitin sequences using the CRISPR Design Tool (Hsu et al., 2013) (See “Key Resources Table” for guide RNA sequences). To construct the *Myc-UBC* cell line, a gene block (Dharmacon) containing the Myc-tag sequence flanked by 1000 bp homology arms on both ends was Gibon-assembled into a SmaI-digested pUC19 vector backbone (Addgene) to make pUC19-Myc-UBC. Single-stranded oligodeoxynucleotides (ssODN, IDT) were designed to introduce the HA-tag and V5-tag at the *RPS27A* and *RPL40* loci, respectively. Plasmid and ssODN donors contained mutations in the PAM sequences at each cut site to prevent Cas9 from cutting the edited loci.

Using a Lonza 4D nucleofector, 2 × 10^5^ HEK 293 cells were nucleofected with preassembled Cas9 RNP complex together with 100 pmol donor ssODN or 750 ng donor plasmid (see “gRNA and Cas9 Preparation” and “Cas9 RNP Nucleofection” above for more details). 48 hours after nucleofection, single cells were dispersed into four 96-well plates to isolate clones. To genotype clones, cells were lysed in QuickExtract^™^ DNA Extraction Solution (Lucigen, see “T7 Endonuclease 1 Assay” for more details), and the edited region was PCR amplified. PCR fragments were TOPO cloned (Thermo Fisher Scientific), and plasmids were analyzed by Sanger sequencing.

### Chemical Genetics

We used a variety of chemical inhibitors to identify the pathways regulating RPS27A and RPL40 depletion. To prevent proteasomal degradation during DNA damage, cells were treated with 50 µM epoxomicin (Calbiochem) for 1 hour before cells were treated with 5 µM etoposide or 50 µM epoxomicin for 16 hours. 10 µM KU 55933 (Tocris) or 10 µM AZ 20 (Tocris) was co-administered with etoposide for 16 hours to inhibit the ATM and ATR pathways, respectively. MDM2-mediated degradation of p53 was prevented after DNA damage with co-administration of 10 µM nutlin (Sigma-Aldrich) and 5 µM etoposide over a 16-hour time course.

To rescue downstream effects of eIF2α phosphorylation after DNA damage, 200 nM ISRIB (Sigma-Aldrich) was added at the same time as etoposide. Cells were treated with 1 µM thapsigargin (Sigma-Aldrich) or 2.5 µM PP242 (Sigma-Aldrich) for 30 minutes as controls for eIF2α phosphorylation or 4E-BP1 hypo-phosphorylation, respectively. To determine the effects of modulating eIF2α phosphorylation on genome editing, HEK-BFP or K562-BFP cells were treated with 10, 50, or 75 µM salubrinal (Tocris) or 200 nM ISRIB (Sigma-Aldrich) for 16 or 24 hours post Cas9 RNP nucleofection.

### Generating RPS27A-SBP Flp-In Cell Lines

RNA from HEK cells was isolated using the DirectZol RNA MiniPrep Kit (Zymo) according to the manufacturer’s protocol. cDNA was generated using SuperScript II Reverse Transcriptase (Thermo Fisher Scientific), and coding regions of RPS27A with and without the N-terminal ubiquitin sequence was PCR amplified and cloned into a pcDNA5/FRT/TO vector backbone (Invitrogen) that had been previously modified to have a constitutive CMV promoter and C-terminal SBP-tag.

To generate stable transgenic cell lines, 1 x 10^6^ HEK Flp-In T-Rex Cells (Invitrogen) were nucleofected using a Lonza 4D nucleofector in according to the Amaxa 4D-Nucleofector^™^ Protocol for HEK293 (Lonza) for large cuvettes with 1.8 µg pOG44 Flp-Recombinase Expression Vector and 0.2 µg pCMV-RPS27A-SBP or pCMV-Ub-RPS27A-SBP. Two days after nucleofection, cells were passaged and placed on media containing 5 µg/ml blasticidin (Invitrogen) and 10 µg/ml Hygromycin B (Thermo Fisher Scientific) until all cells from a control plate nucleofected with pmaxGFP(tm) Vector (Lonza) were dead. Flp-In cell lines were validated using anti-SBP Westerns and Sanger sequencing of the transgenic insert.

### Ubiquitin Blots of Denatured RPS27A-SBP

RPS27A-SBP Flp-In HEK cells were lysed in binding buffer (300 mM NaCl, 0.5% NP-40, 50 mM Na_2_HPO_4_, 50 mM Tris pH 8) with 8M urea using the protocol detailed in “Western Blotting.” Samples were diluted 1:3 with binding buffer, and normalized lysates were incubated at 4 °C for 30 minutes with 60 µl buffer-equilibrated Dynabeads^™^ M-270 Streptavidin (Invitrogen). Beads were washed 5 times with 200-500 µl binding buffer containing 1M NaCl. To elution proteins, beads were boiled in 25 µl 1X NuPAGE loading buffer at 97 °C for 5 minutes. Westerns were performed as detailed in “Western Blotting.”

### siRNA Knockdowns

siRNA oligonucleotides (see “Key Resources Table” below) were transiently transfected into cells using RNAiMAX (Invitrogen) according to manufacturer instructions. For each well in 12-well plate, 120 pmol siRNA and 3.6 µl RNAiMAX were used. Cells were transfected with siRNAs 24 hours prior to drug treatment or Cas9 nucleofection.

### pHA-Ub Immunoprecipitations

10 cm plates HEK293 or RPS27A-SBP Flp-In cells were transiently transfected with 10 µg of HA-UB plasmid (gift from Rape lab) with Lipofectamine 3000 (Thermo Fisher Scientific) for 48 hours. Immunoprecipitation was performed using Pierce Anti-HA Magnetic Beads Kit (Thermo Fisher) according to the manufacturer. 1 mg of cell lysate and 50 µl beads were used to perform each immunoprecipitation. After overnight incubation at 4 °C, the beads were washed twice with IP buffer supplemented with 500 mM NaCl and twice with regular IP buffer and proteins were eluted by boiling samples at 98°C in 1X NuPAGE LDS sample buffer (Thermo Fisher) for 5 min. When siRNA was used, cells were first transfected with siRNAs and after 24 hours, with the HA-Ub plasmid. Lysates were prepared 48 hours after the second transfection with drug treatment with epoxomicin and etoposide occuring 17 hours and 16 hours before lysis, respectively.

### Bulk Translation Assays

10 cm plates of HEK cells were washed with PBS then placed in 25 µM Click-IT L-Azidohomoalanine (Thermo Fisher Scientific) in DMEM, high glucose, no glutamine, no methionine, no cysteine (Gibco) with 10% FBS for 2 hours. Cells were trypsinized then pelleted at 400 x g for 5 minutes. Cells were washed three times with PBS before being lysed in 200 µl lysis buffer (1% SDS, 50 mM Tris HCl, pH 8.0, 1X Halt Protease Inhibitor Cocktail, Thermo Scientific) with 150 U/ml benzonase nuclease to digest DNA and RNA. Lysates were incubated for 30 minutes on ice, vortexed for 5 seconds, and spun at 18,000 x *g* for 10 minutes at 4 °C. Protein content of the supernatants was normalized using the Pierce BCA Protein Assay (Thermo Fisher Scientific). 1 µl 10 mM IRDye 800CW DBCO Infrared Dye was added to the lysates, and the lysates were incubated for 2 hours at RT. Unbound IR Dye was removed using a Zeba Column, 7K MWCO, 0.5 mL (Thermo Fisher Scientific). For dot blot analysis, a Bio-Dot Microfiltration Apparatus (Bio-Rad) was used according to the manufacturer’s protocol and 20 µl sample dilutions were added to wells. Membranes were imaged on a LI-COR Odyssey CLx Imager. For protein gel analysis, lysates were combined with 2X Laemmli Buffer, incubated at 97 °C for 5 min, then run on a Nupage 4-12% Bis-Tris Gel (Invitrogen) at 200V fro 40 min. The gel was washed with PBS (3 x 5 minutes) before imaging with a LI-COR Odyssey CLx Imager.

### NGS Analysis of Editing Outcomes

HEK cells carrying a single copy of a BFP transgene were nucleofected with Cas9-sgBFP or dCas9-sgBFP and recovered in media containing 75 µM salubrinal or 200 nM ISRIB for 24 hours. gDNA extraction and 50 µl PCRs (PCR1, see “Key Resources Table” for sequences) of the edited genomic loci were prepared as detailed in “T7 Endonuclease 1 Assay.”

PCR1 reactions were cleaned up using SPRI bead purification. A 50 mL stock solution of SPRI beads was prepared in advance: 1 ml SPRI beads (Sera-Mag SpeedBeads® Carboxyl Magnetic Beads) were brought to room temperature and washed three times with TE buffer before suspended to 50 ml in 18% PEG-8000, 1 M NaCl, 10 mM Tris-Cl (pH 8.0), 1 mM EDTA, and 0.055% Tween-20. To purify PCR products, 90 µl SPRI bead suspension solution was added to 50 µl PCR reactions in a 96 well plate. The solution was mixed 10 times with a pipette and incubated at room temperature for 1 minute. The plates were placed on a magnetic stand for 2 minutes, and the supernatant was discarded. 200 µl 80% ethanol was added then removed after 2 minutes while the plate remained on the magnetic stand. The ethanol wash and removal steps were repeated one more time for a total of two washes. The beads were left to air dry for 3-10 minutes. To elute the purified PCR1 products from the beads, beads were resuspended in 20 µl ultra-pure water and incubated for 2 minutes. The plate was placed on a magnetic stand for 1 minute, and the supernatant was collected. Concentrations of purified PCR1 products were quantified using the Qubit^™^1X dsDNA HS Assay with the Invitrogen Qubit^™^4 Fluorometer (Thermo Fisher Scientific) as per the manufacturer’s instructions.

To add Illumina adaptors to the PCR1 products, a second PCR reaction was performed with PrimeSTAR GXL DNA Polymerase (Takara) in a 25 µl reaction with 10 ng PCR1 product and 0.5 µM adaptor according to the manufacturer’s instructions. We used adaptors from a custom set of 960 unique combinatorial Illumina TruSeq indices (IDT) supplied by the Vincent J. Coates Genomics Sequencing Laboratory at UC Berkeley. The samples were amplified for 12 cycles consisting of: 95 °C for 10 seconds, 57 °C for 15 seconds, and 65 °C for 30 seconds. PCR2 products were purified and quantified as detailed above. A Biomek FXp Liquid Handler (Beckman Coulter) was used to pool 50 ng of each PCR product, and a 5300 Fragment Analyzer (Advanced Analytical) was used to assess the concentration and quality of the pool before sequencing.

Samples were deep sequenced on an Illumina MiSeq at 300 bp paired-end reads to a depth of at least 10,000 reads. A modified version of CRISPResso (Pinello et al., 2016) was used to analyze editing outcomes and to plot mutation position distributions. Briefly, reads were adapter trimmed then joined before performing a global alignment between sequence reads and the *BFP* reference sequences using NEEDLE (Li et al., 2015). Indel rates were calculated as any reads where an insertion or deletion overlaps the cut site or occurs within three base pairs of either side of the cut site divided by the total number of reads.

### Ribosome Profiling and RNA-seq

Paired ribosome profiling and RNA-seq experiments were conducted on HEK 293 cells lysed 36 and 72 hours after Cas9 or dCas9 RNP nucleofection. Cas9 and dCas9 complexed with sgIntron, a guide targeting intron 12 of *JAK2,* were nucleofected using the protocols detailed in “Cas9 RNP Nucleofections” above. Four small-scale nucleofections were pooled directly into one 10 cm plate to create one biological replicate with each experimental condition having two biological replicates. Due to recent reports about IVT guide RNAs inducing interferon responses in cells (Kim et al., 2018; Wienert et al., 2018), synthetic gRNAs (Synthego) were used at the 36 hour time point.

Ribosome profiling was conducted as detailed in (McGlincy and Ingolia, 2017) with the following modifications. Since Epicentre discontinued the yeast 5′-deadenylase (Cat# DA11101K) we used in our published protocol, we cloned a 5′-deadenylase (*HNT3*) from the thermotolerant yeast *Kluyveromyces marxianus* into the pET His6 TEV LIC cloning vector (2B-T) backbone (gift from Scott Gradia to Addgene). Recombinant 6xHis-TEV-Km-HNT3 was purified from *E. coli* using a Nickel column purification (HisTrap FF Crude column, GE Life Sciences). Protein eluted from the column with imidazole was cleaved with TEV protease, and the residual His tag was removed using a Nickel column. The recombinant protein subsequently purified using size exclusion chromatography (Sephacryl S-300 16/60 column, GE Life Sciences). 0.5 µl of purified protein was added in place of the yeast 5′-deadenylase during ribosome profiling, and the reaction was incubated at 37 °C instead of 30 °C.

We also deviated from the McGlincy and Ingolia 2017 protocol by using CircLigase I instead of CircLigase II. We made this change after concerns about the nucleotide bias of CircLigase II were reported in (Tunney et al., 2018). Therefore, we reverted to using CircLigase I as previously detailed in (Ingolia et al., 2012) with a 2 hour incubation step.

Total RNA for mRNA-seq was isolated from 50 µl cell lysate using the DirectZol^™^ RNA MiniPrep Kit (Zymo) according to the manufacturer’s protocol. Sequencing libraries were prepared using the TruSeq Stranded Total RNA Library Kit with Ribo-Zero Gold (Illumina). Ribosome profiling and RNA-seq libraries were sequenced as 50 nt single-end reads on an Illumina HiSeq 4000.

Reads from ribosome profiling were processed as detailed in (McGlincy and Ingolia, 2017). Ribosome profiling and RNA-seq reads from the 36 hour time point were aligned with HiSat2 (Kim et al., 2015) to the Human GENCODE Gene Release GRCh38.p2 (release 22); reads from the 72 hour time point were aligned with TopHat (Trapnell et al., 2009) to GRCH38.p7 (release 25). Alignments were indexed using Samtools (Li et al., 2009), and the number of reads per transcript was tabulated using fp-count (Ingolia et al., 2014) with the basic gene annotations from GRC38.p2 (36 hr) and GRCh38.p7 (72 hr). Differential changes in gene expression were calculated using DESeq2 (Love et al., 2014) with a cutoff of FDR < 0.1 for per-gene significance. Translational efficiency (the ratio of ribosome footprints to mRNA-seq transcripts) calculations and significance tests were made in DESeq2 using a design matrix that tests the ratio of ratios (design = ∼ A + B + A:B, where A is Cas9 type and B is library type) with FDR < 0.1.

Cumulative distribution functions and Mann-Whitney-Wilcoxon tests with ribosome profiling and RNA-seq data were calculated in RStudio. Three gene lists were used for this analysis: ISR targets, ribosome proteins, and DSB break repair genes. ISR (Integrated Stress Response) targets are the 78 genes identified by (Sidrauski et al., 2015) to have a statistically significant, greater than twofold change in translational efficiency after tunicamycin treatment. (6 of the 78 genes were removed from analysis because we were unable to identify corresponding GRCh38 Ensembl gene IDs from the original GRCh37 UCSC gene IDs listed in Sidrauski et al., 2015.) Ribosome proteins are the 80 core ribosomal protein genes expressed in humans. DSB break repair genes are 44 genes from the union of genes annotated as DSB repair genes in **Table S3** from (Chae et al., 2016) and on the University of Pittsburgh Cancer Institute’s DNA Repair Database website (https://dnapittcrew.upmc.com/db/index.php).

## QUANTIFICATION AND STATISTICAL ANALYSIS

Bar graphs, scatterplots, stripcharts, and cumulative distribution function plots were created with RStudio version 1.0.136 running R version 3.3.2. Standard statistical analyses such as standard deviation calculations and Mann-Whitney-Wilcoxon tests were conducted in R. FDR values for RNA-seq and ribosome profiling were calculated using the Wald test in DESeq2 as described in (Love et al., 2014). Statistical details of experiments such as sample size (n) can be found in the figures and figure legends. For this paper, n = number of biological replicates and SD = standard deviation assuming a normal distribution.

## DATA AND SOFTWARE AVAILABILITY

Ribosome profiling and mRNA-Seq data are available from NCBI GEO, Accession #GSE122615.

## KEY RESOURCES TABLE

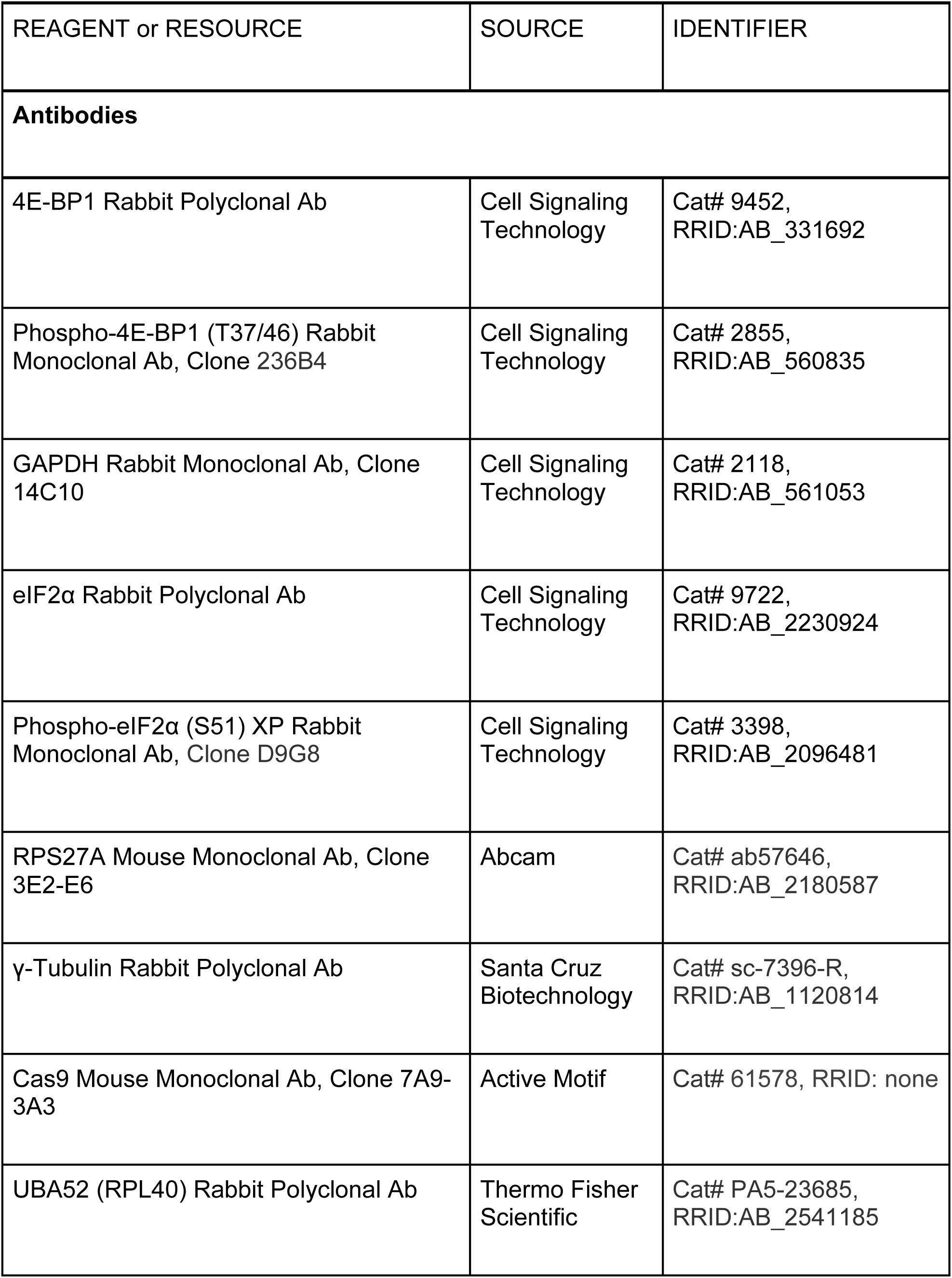
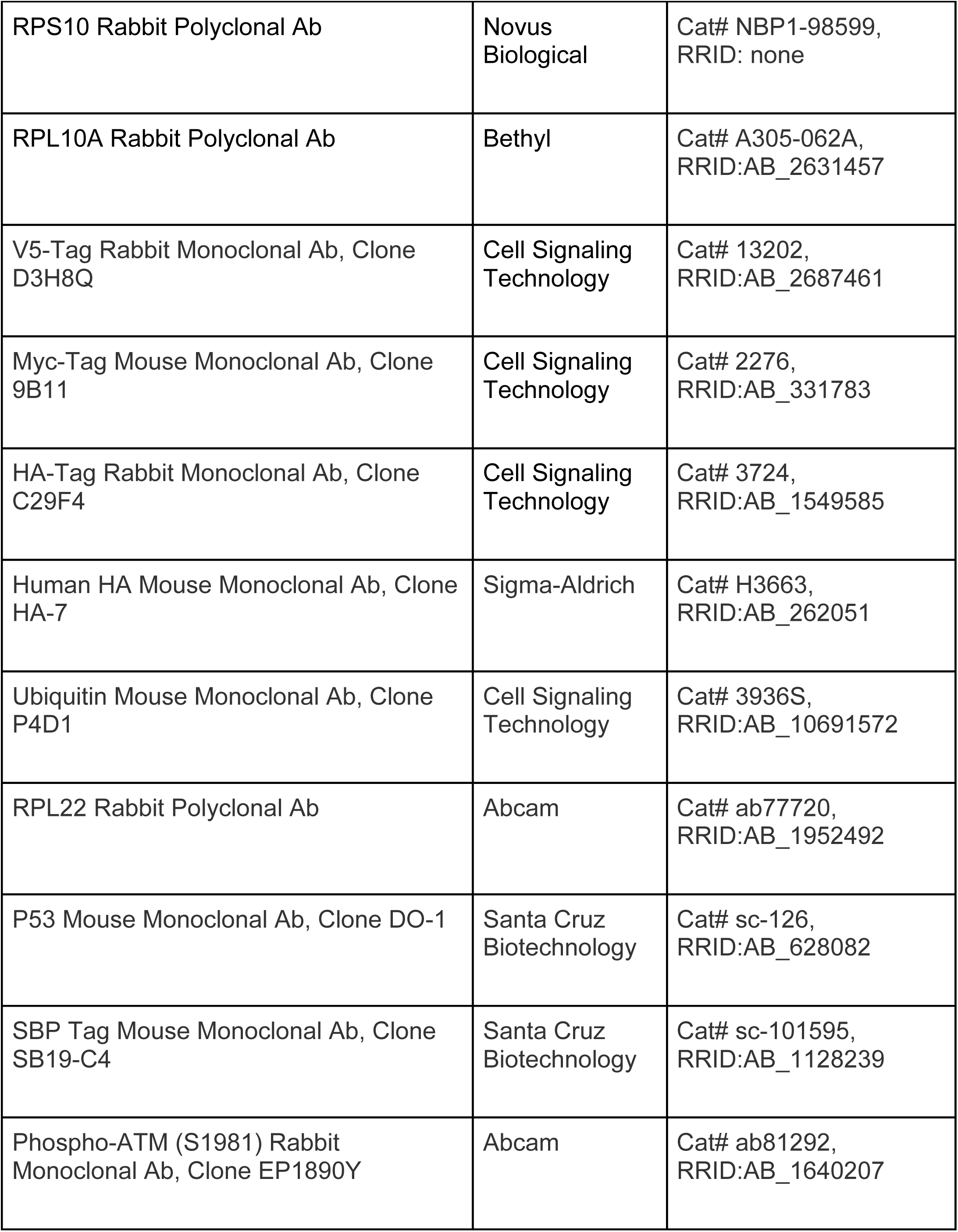
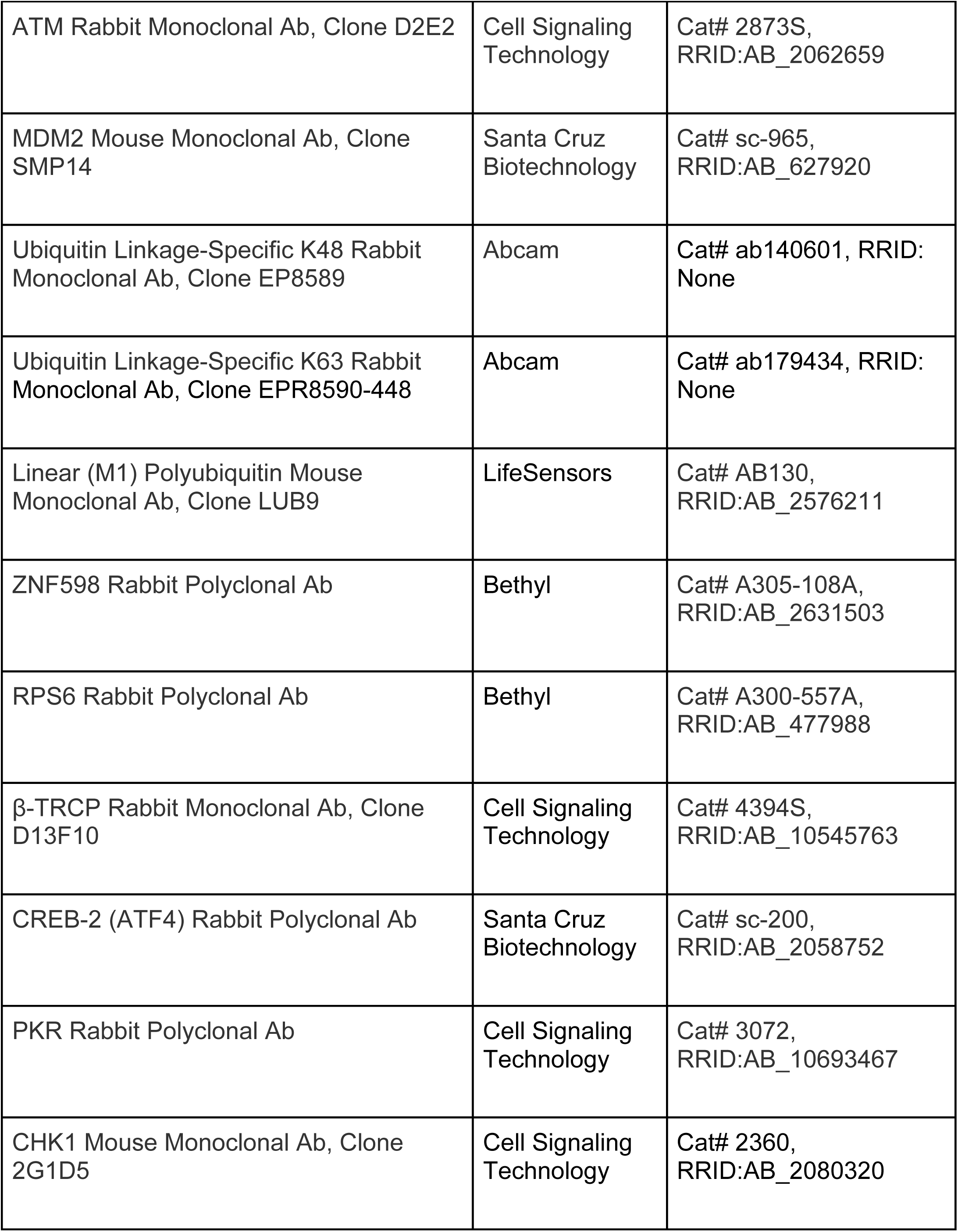
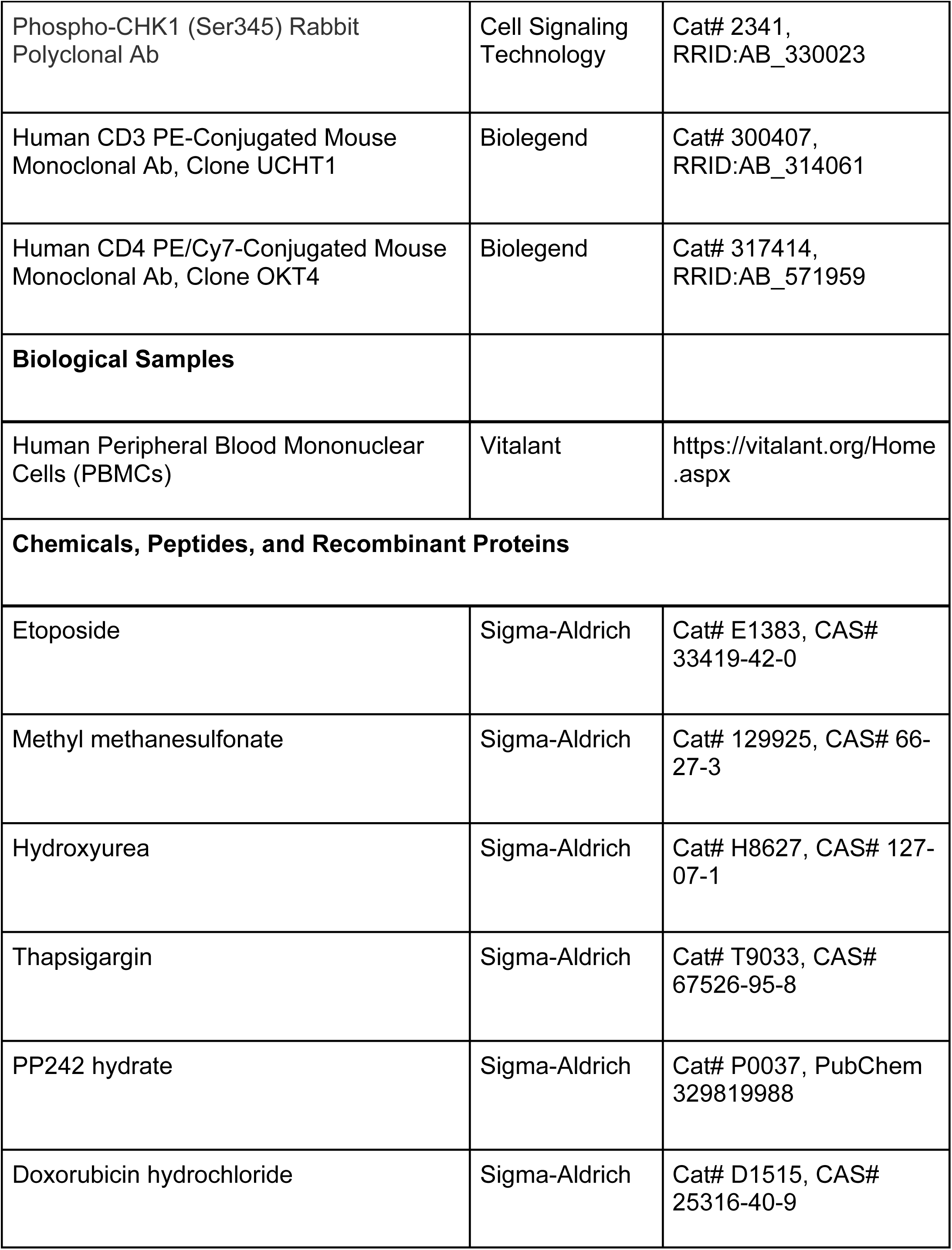
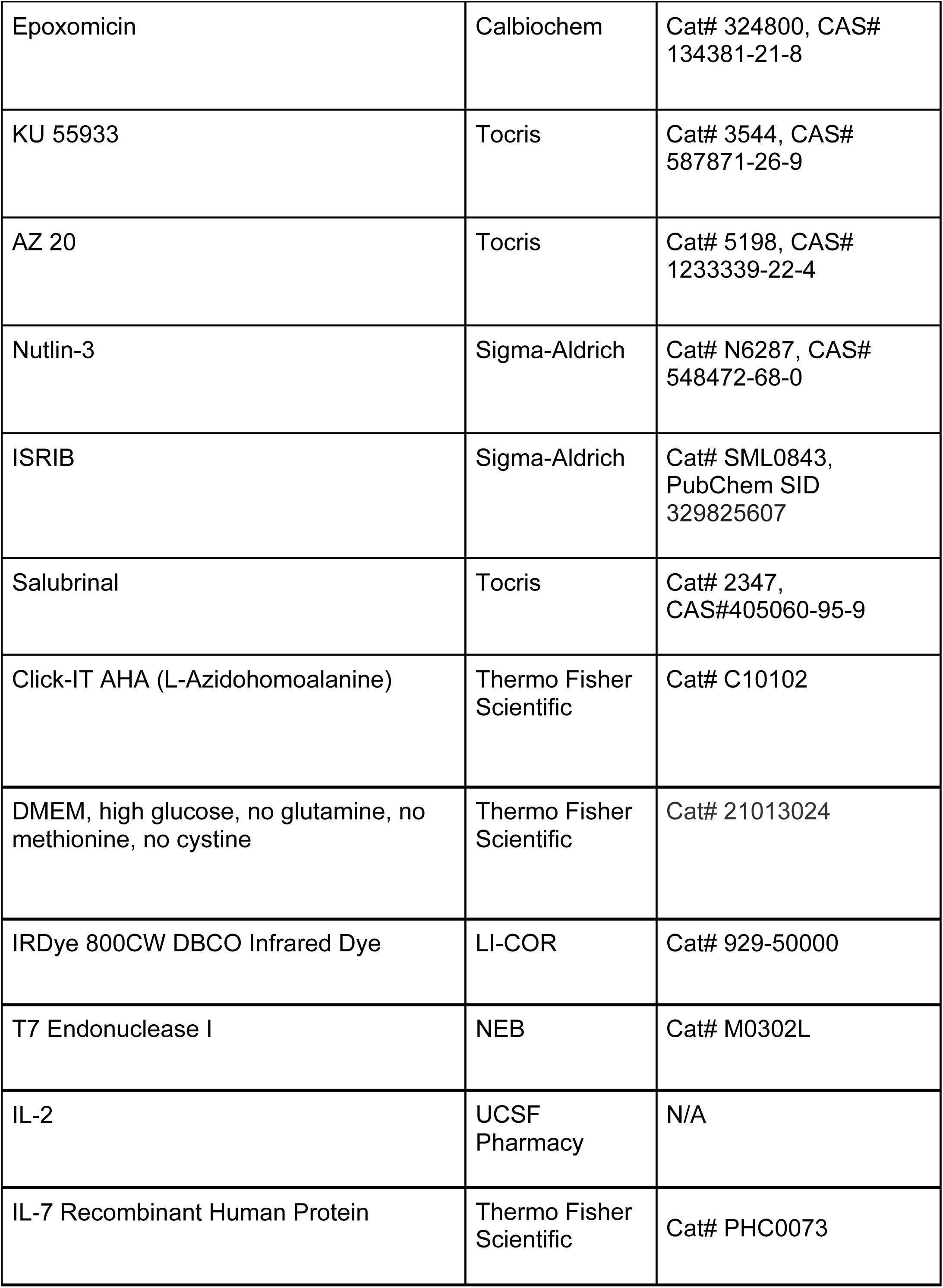
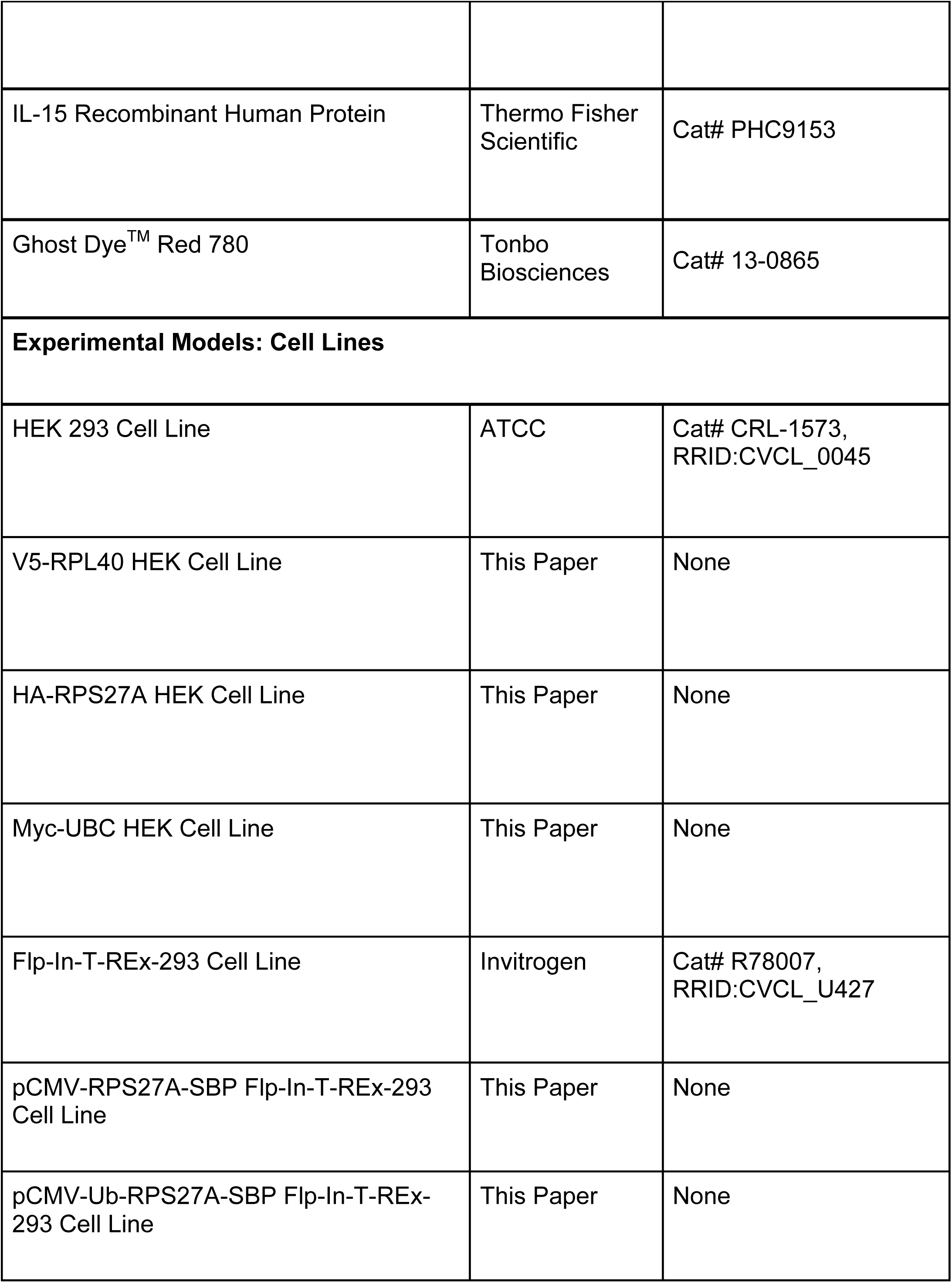
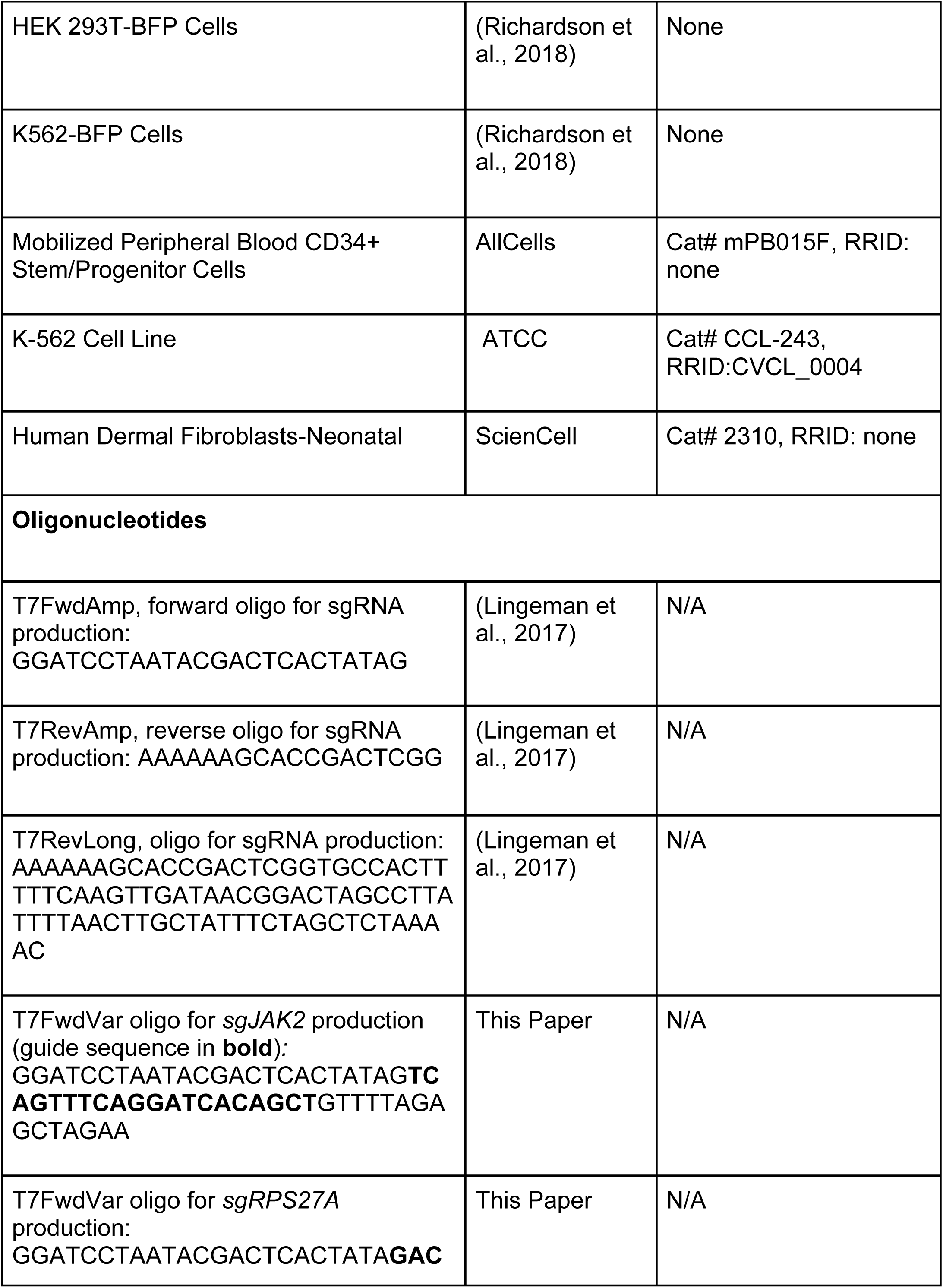
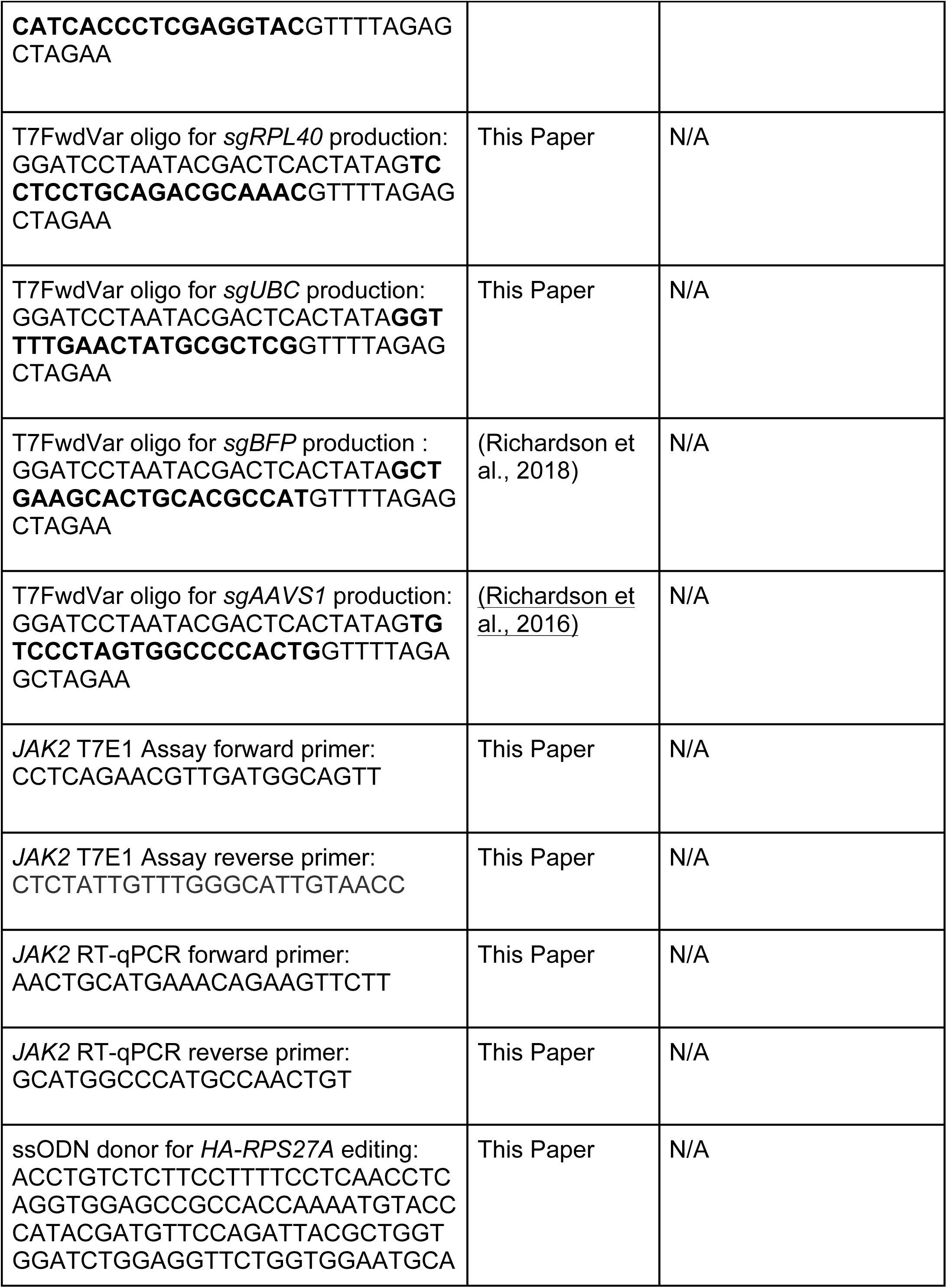
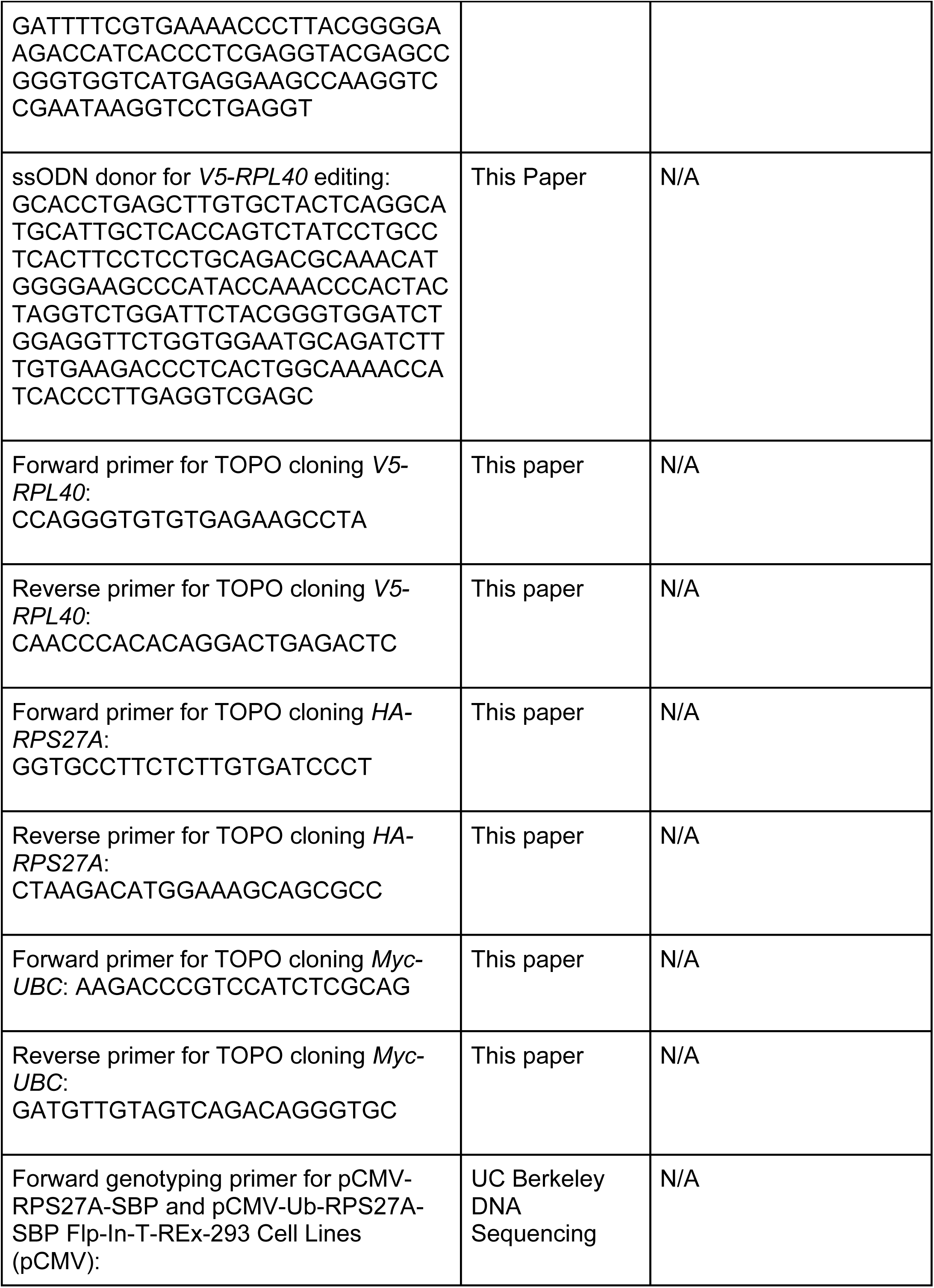
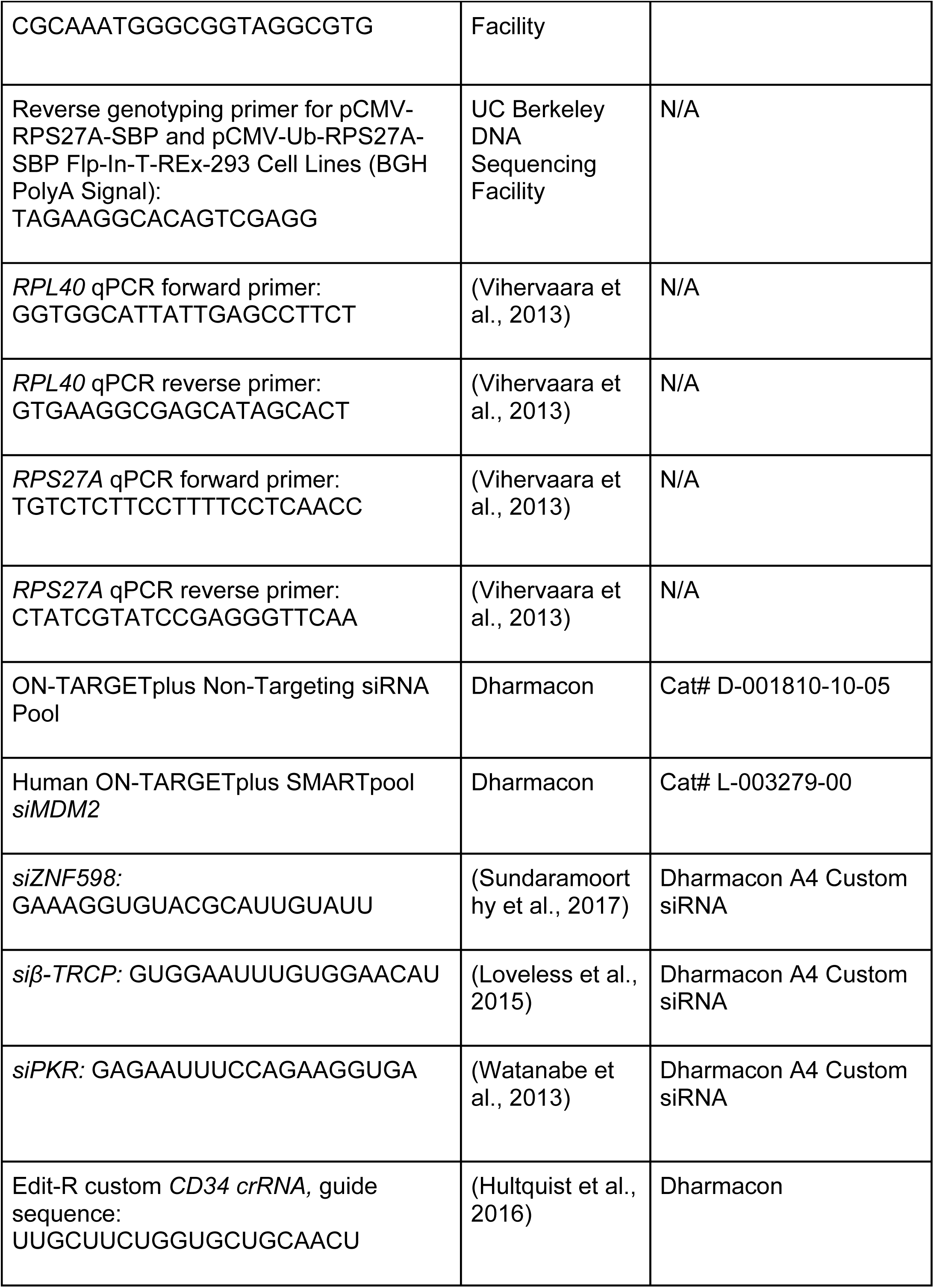
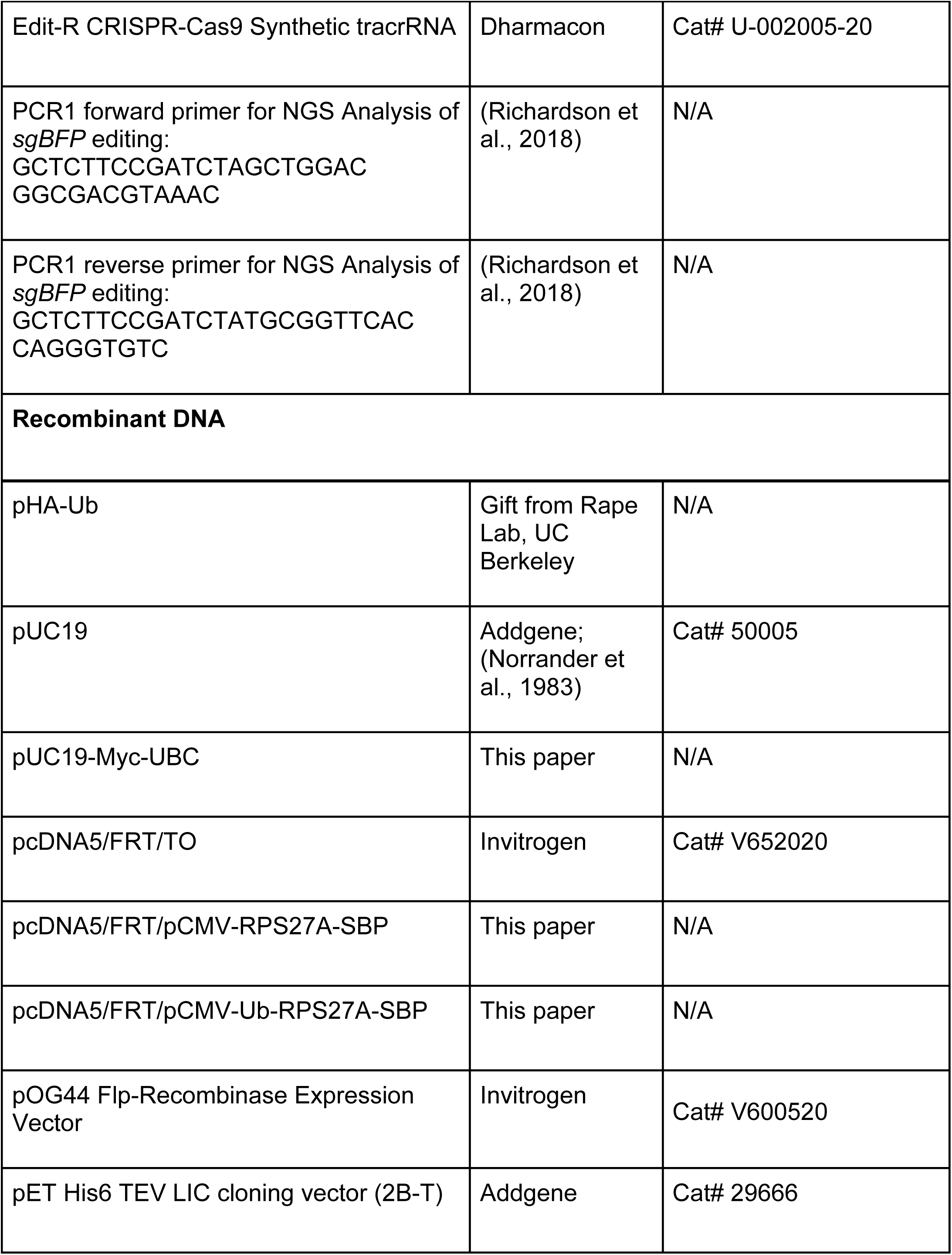
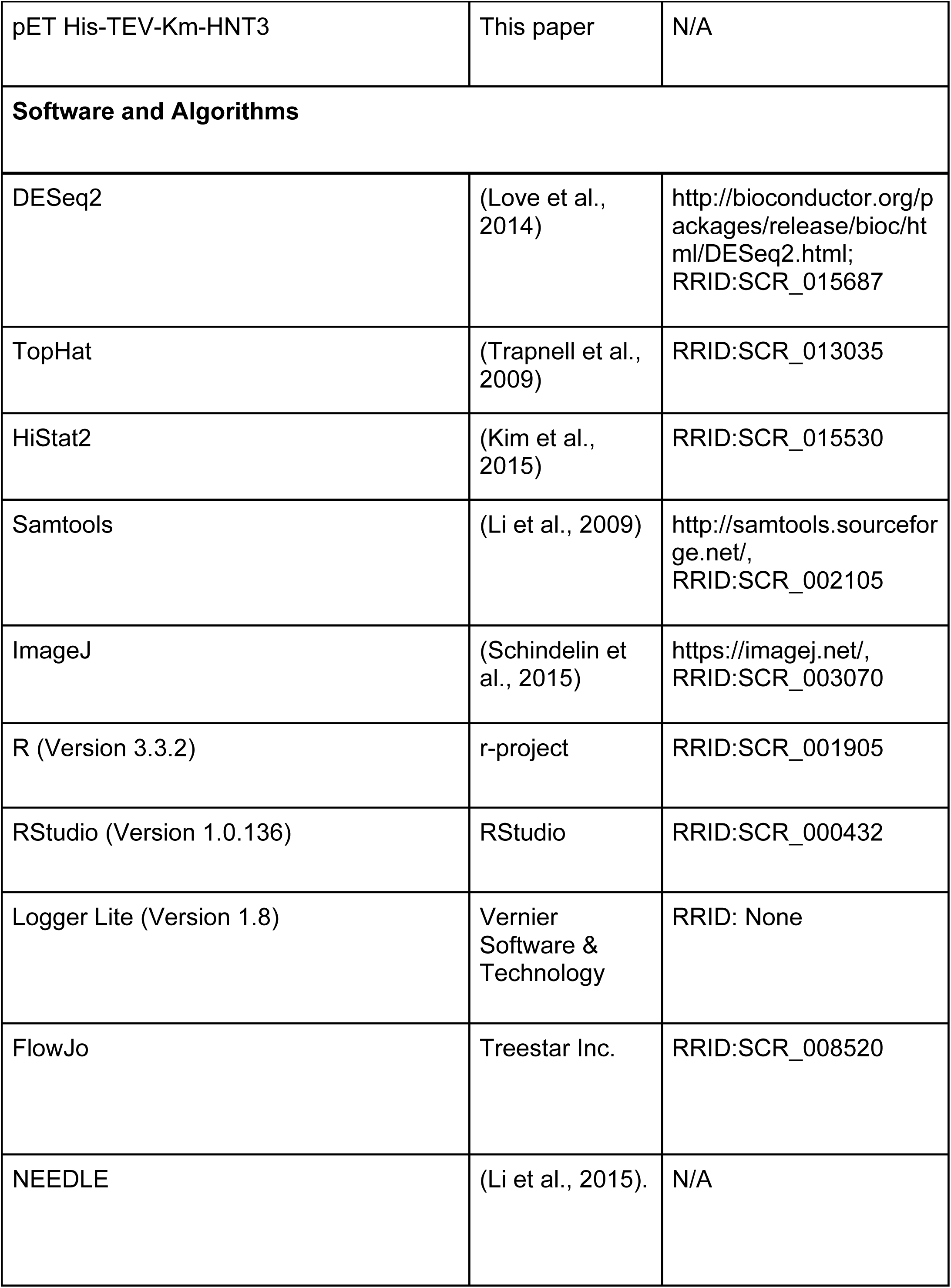
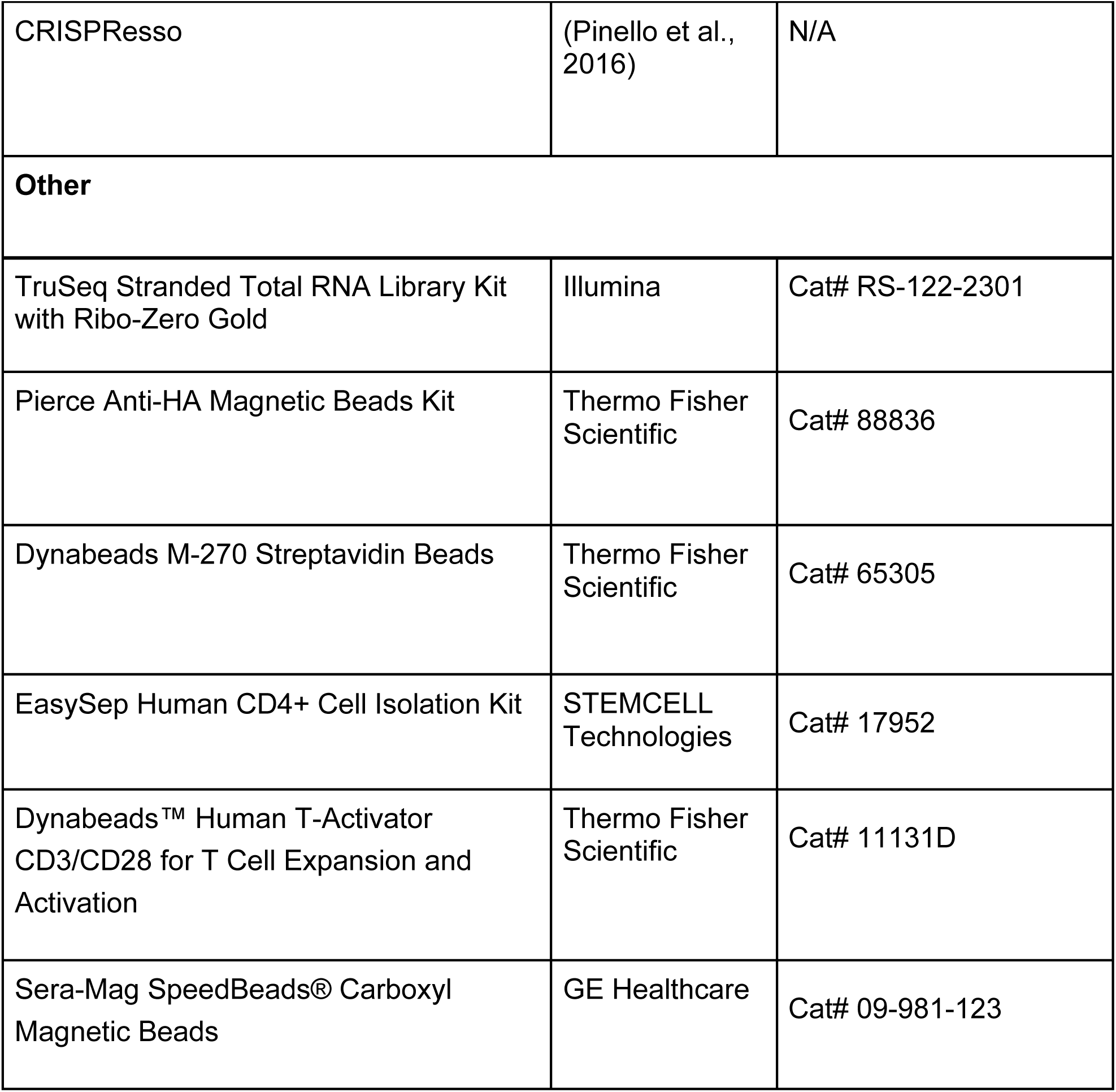

## Supplemental Information

**Table S1: NGS Allele Frequency Analysis (rel to Fig. 5)**

Sheet 1: Summary of Alignments and Indel Frequencies
Sheets 2-12: Allele Frequencies per NGS Sample

**Table S2: Ribosome Profiling and RNA-seq DESeq2 Analysis (rel to Fig. 6)**

Sheet 1: Ribosome Profiling, 36 Hours
Sheet 2: RNA-seq, 36 Hours
Sheet 3: Translational Efficiency, 36 Hours
Sheet 4: Ribosome Profiling, 72 Hours
Sheet 5: RNA-seq, 72 Hours
Sheet 6: Translational Efficiency, 72 Hours

**Table S3: Target Gene Lists for CDF Plots (rel to Fig. 6 and Fig. S6)**

Sheet 1: Integrated Stress Response (ISR) Genes, (Sidrauski et al., 2015)
Sheet 2: Ribosome Protein Genes
Sheet 3: DSB Repair Genes, union of genes annotated as DSB repair genes from (Chae et al., 2016) and University of Pittsburgh Cancer Institute’s DNA Repair Database
Sheets 4-21: DESeq2 results for target genes that were used to generate Figures 6E-H and S6C. These gene lists represent the intersection of the target gene lists and all genes identified at 36 and 72 hours.

## Supplemental Figure Legends

**Figure S1.**
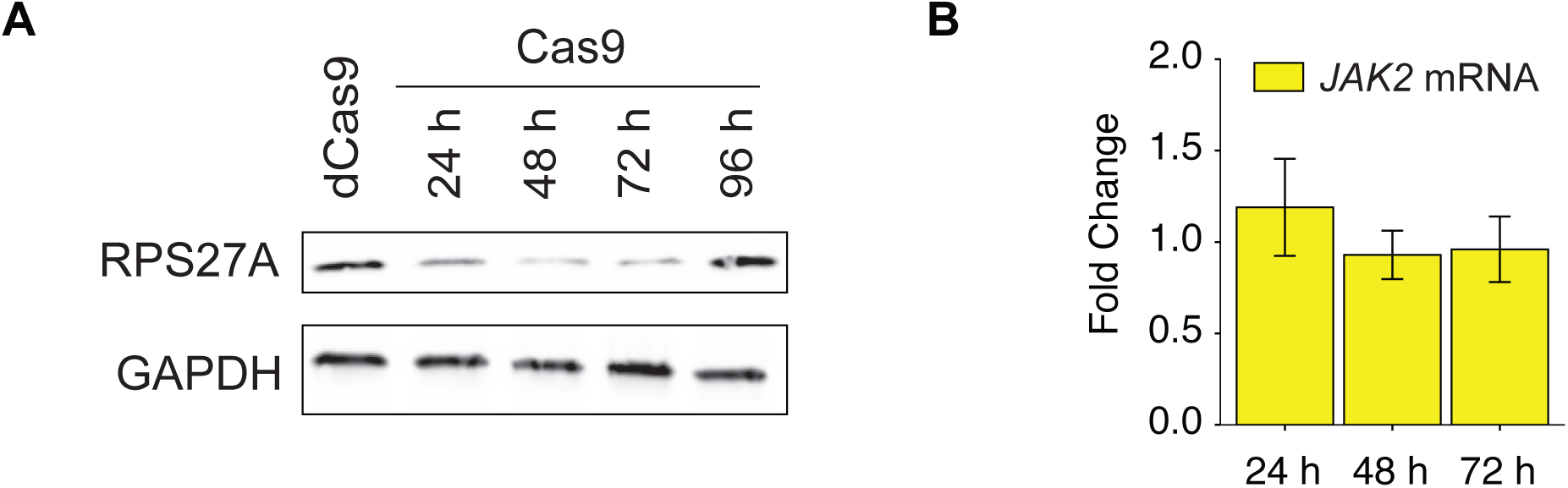
(Related to Fig. 1). Ribosome proteins RPS27A and RPL40 are downregulated after genome editing with Cas9. A. As in Figure 1A, showing recovery of RPS27A at 96 hours post-nucleofection.
B. Genome editing does not affect *JAK2* mRNA abundance. Fold changes were calculated using the 2^-ΔΔCt^ method with Cas9 without sgIntron (apo Cas9) as the control and *GAPDH* as the reference gene (n = 3, error bars = SD).

**Figure S2.**
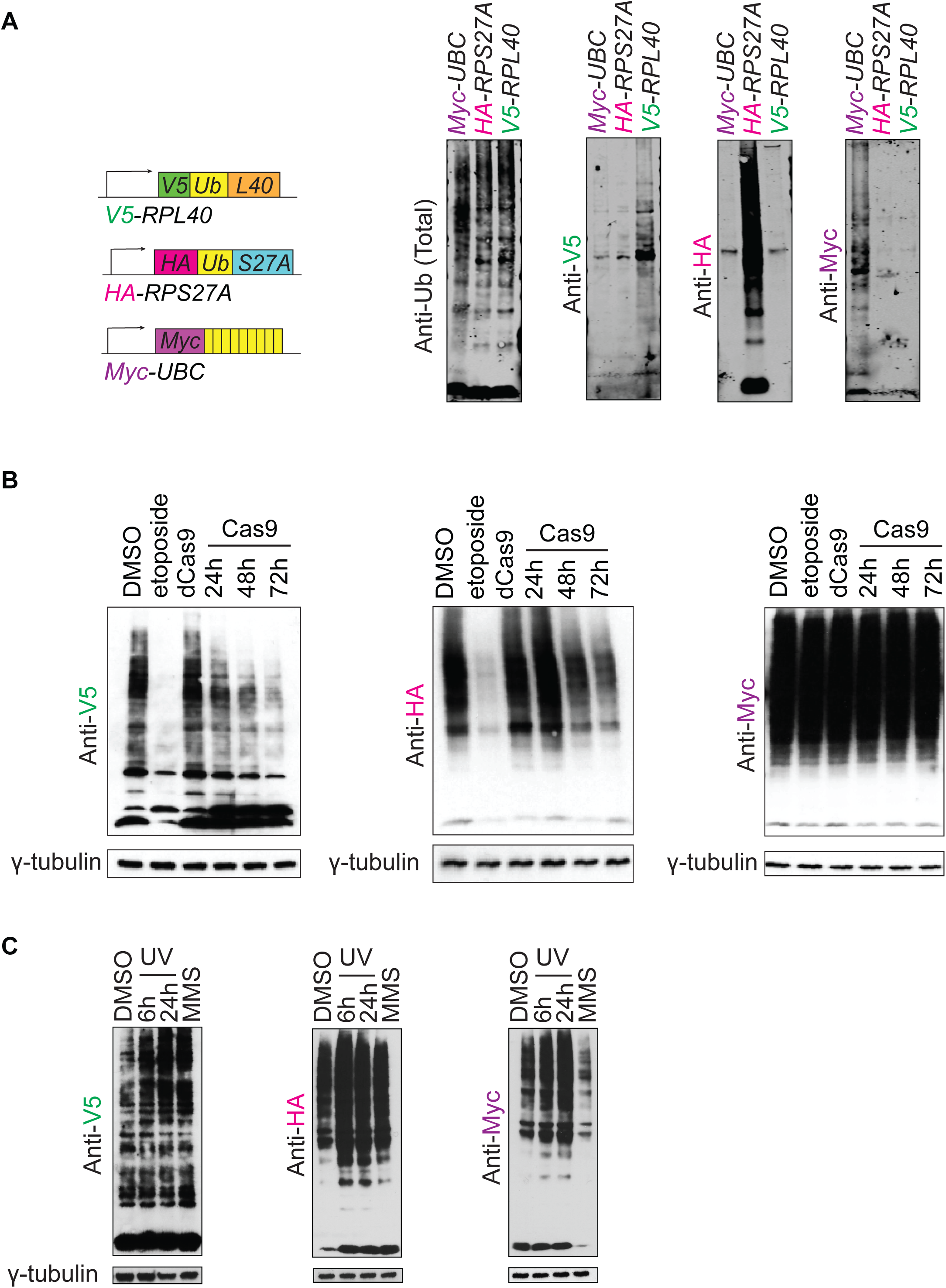
(Related to Fig. 2). Ubiquitins translated from *RPS27A* and *RPL40* decrease after dsDNA breaks. A. Western blotting of HEK 293 cell lines edited to introduce epitope tags at the endogenous *RPL40, RPS27A*, and *UBC* loci.
B. As in **Figure 2C**, nucelofection with dCas9 RNPs (72 hours) does not lead to depletion of V5-Ub and HA-Ub, demonstrating that their depletion is due to Cas9 DSBs.
C. Tagged ubiquitin expression in edited HEK cells after UV radiation (20 J/m^2^) or treatment with 0.03% MMS for 1 hour.

**Figure S3.**
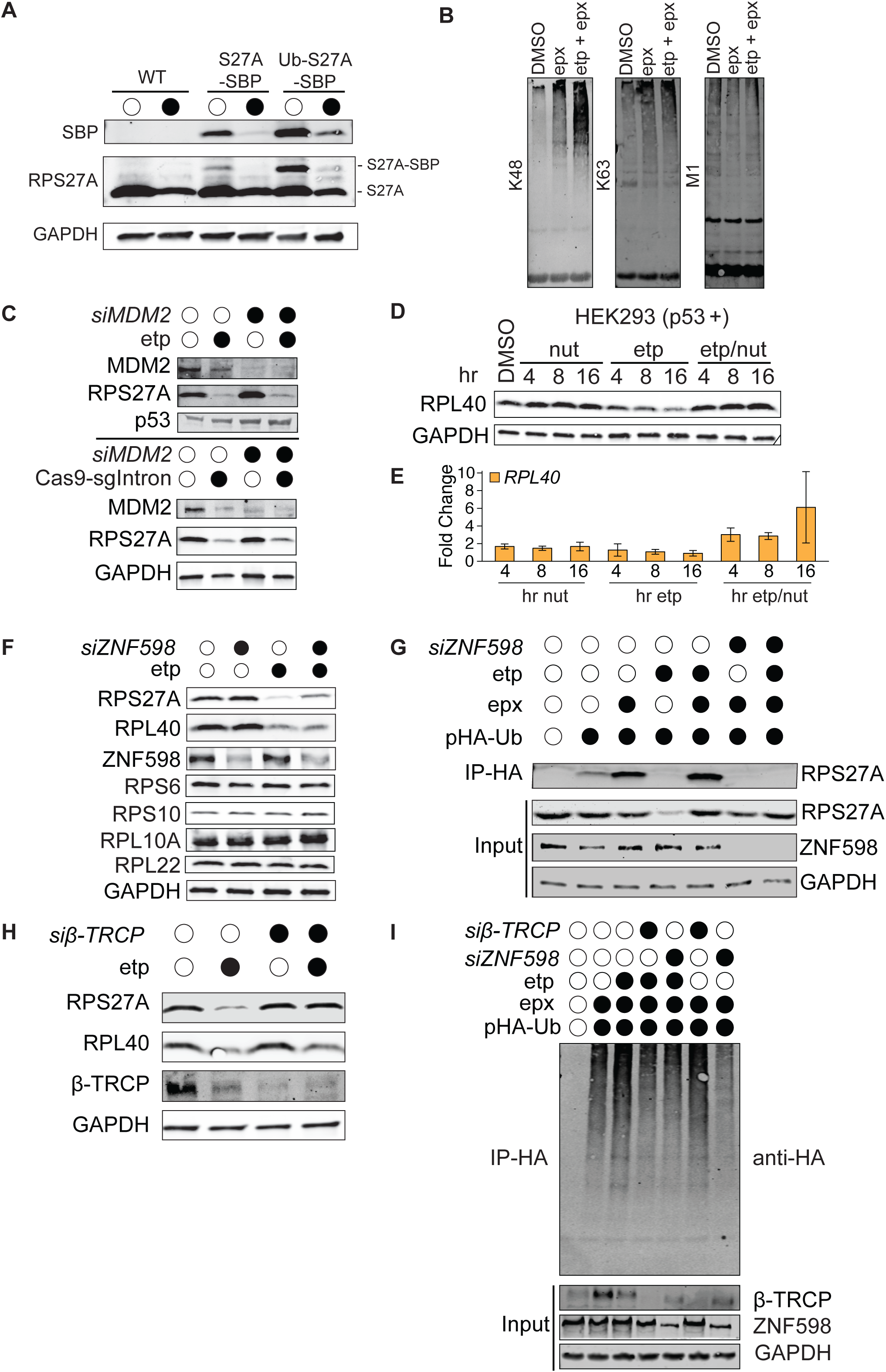
(Related to Fig. 3). RPS27A is proteasomally degraded after dsDNA damage. A. Western blotting of RPS27A and SBP tag in HEK Flp-In cell lines with stable, single-copies of *pCMV-Ub-RPS27A-SBP* or *pCMV-RPS27A-SBP* transgenes after 5 µM etoposide or DMSO treatment for 16 hours. Note that *RPS27A* transgenes lack the endogenous promoter and UTR sequences.
B. Western blotting for K48, K63, and M1 ubiquitin linkages on affinity-purified RPS27A-SBP indicates constitutive and etoposide-induced K48-linked ubiquitin chains. HEK Flp-In cell lines expressing *pCMV-RPS27A-SBP* were treated with epoxomicin (50 µM, 17 hours) and etoposide (5 µM, 16 hours).
C. Western blotting demonstrates RPS27A depletion is insensitive to *MDM2* knock-down 16 hours after 5 µM etoposide treatment or 72 hours after Cas9-sgIntron nucleofection. Non-targeting siRNAs, DMSO, and Cas9 without a guide served as negative controls.
D. Western blotting of RPL40 shows that nutlin (10 µM) treatment rescues RPL40 depletion induced by etoposide (5 µM) in HEK cells.
E. Abundance of *RPL40* transcripts increases after co-administration of nutlin and etoposide.
F. Western blotting to monitor RPS27A and RPL40 after *ZNF598* knock-down relative to a non-targeting control siRNA.
G. Western blotting against RPS27A protein following anti-HA immunoprecipitations from HEK cell lysates transfected with a plasmid expressing HA-Ub. Cells were transfected with siRNAs and, after 24 hours, with the HA-Ub plasmid. Lysates were prepared 48 hours after the second transfection.
H. Western blotting against RPS27A protein following siRNA knockdown of *β-TRCP* (or a non-targeting control siRNA) and etoposide treatment.
I. As in (G), using an HEK cell line expressing a single-copy *pCMV-RPS27A-SBP* transgene.

**Figure S4.**
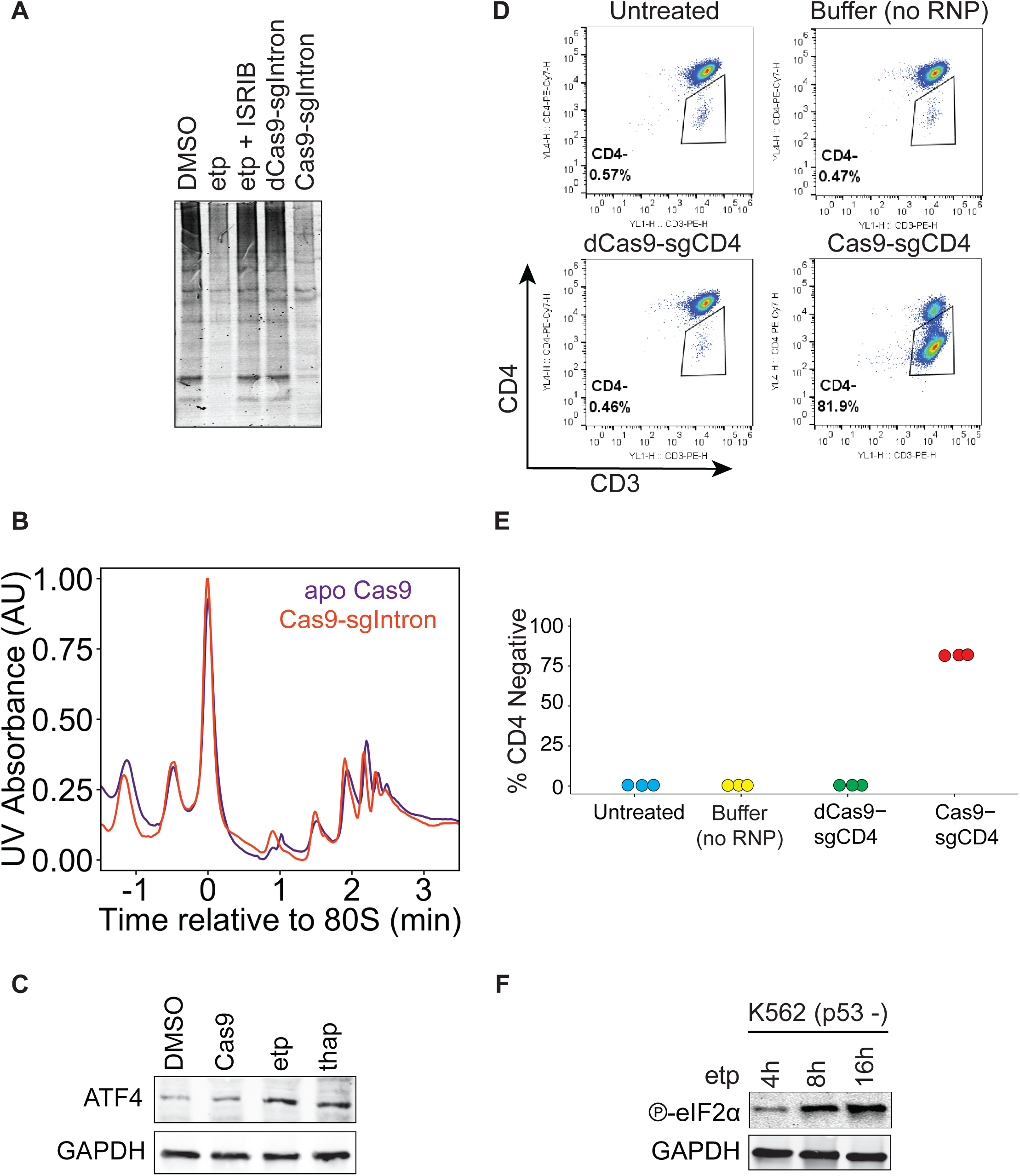
(Related to Fig. 4) Double-strand DNA breaks lead to eIF2α phosphorylation and reduced translation initiation. A. IR800 LI-COR-image of SDS-PAGE gel with L-AHA labeled lysates depicted in (**Figure 4H**).
B. Polysome profiles of HEK cells 72 hours after nucleofection with active Cas9-sgIntron RNP, or Cas9 without guide (apo Cas9).
C. Western blotting of ATF4 induction. Cells were harvested 72 hours after nucleofection with Cas9-sgIntron or 16 hours after treatment with 5 µM etoposide. Cells treated with DMSO for 16 hours or 1 µM thapsigargin for 30 minutes served as negative and positive controls respectively.
D. Examples of flow cytometry editing efficiency analysis of T-cells nucleofected with Cas9-sgCD4 in tandem with cells depicted in **Figure 4F**. T-cells were stained with anti-CD3-PE, anti-CD4-PE-Cy7, and GhostDye780 (to mark dead cells).
E. Average percentage of edited, CD4 negative T-cells three days after Cas9-sgCD4 electroporation as determined by FACS (n = 3).
F. Western blotting of eIF2α (S51) phosphorylationin K562 cells treated with 5 µM etoposide.

**Figure S5.**
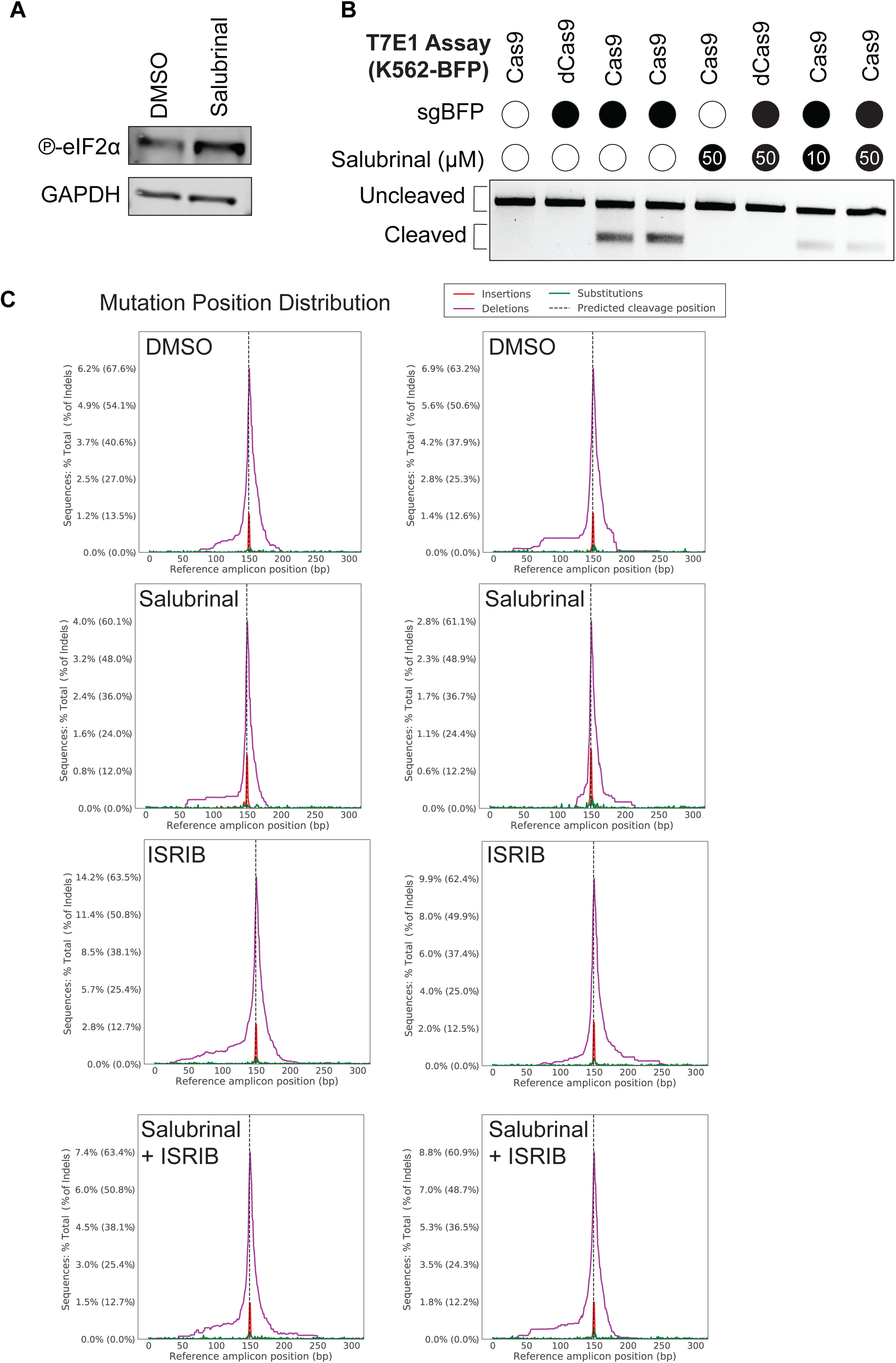
(related to Fig. 5). Modulating eIF2α phosphorylation alters genome editing outcomes. A. Western blotting of eIF2α (Ser51) phosphorylation in HEK-BFP cells treated with 75 µM salubrinal or DMSO for 24 hours.
B. T7 Endonuclease 1 cleavage assay. K562-BFP cells were nucleofected with sgBFP-Cas9 (or dCas9) RNPs and treated with 10 or 50 µM salubrinal for 16 hours.
C. Mutation distribution plots of NGS reads with insertions or deletions (% reads with indels) from gDNA PCRs of HEK-BFP cells nucleofected with sgBFP-Cas9 (or dCas9) RNPs and treated with 75 µM salubrinal or 200 nM ISRIB for 24 hours.

**Figure S6.**
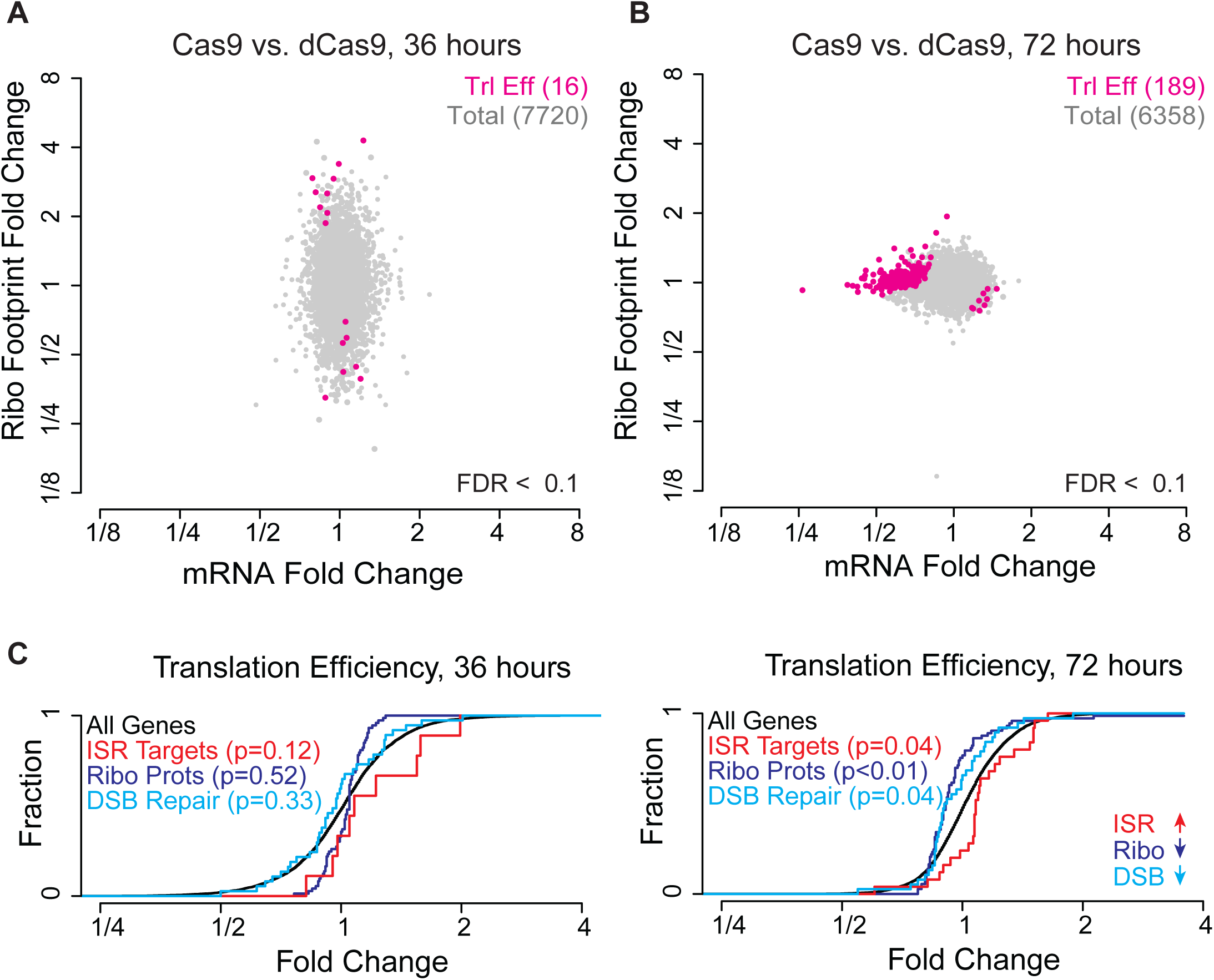
(Related to Fig.6). Genome editing initiates a translational response that proceeds long-term transcriptional changes. A. As in **Figure 6C**, with pink marking genes with significant changes in translation efficiency, the ratio of ribosome footprints to mRNA transcripts (Wald test, FDR adjusted *p*-value < 0.1).
B. As in (A) for 72-hour ribosome profiling and RNA sequencing data.
C. As in Figures 6E through 6H, for translation efficiency. See **Table S3** for target gene lists.

